# Low doses of the organic insecticide spinosad trigger lysosomal defects, ROS driven lipid dysregulation and neurodegeneration in flies

**DOI:** 10.1101/2021.02.03.429672

**Authors:** Felipe Martelli, Zuo Zhongyuan, Julia Wang, Ching-On Wong, Nicholas E. Karagas, Ute Roessner, Thusitha Rupasinghe, Kartik Venkatachalam, Trent Perry, Philip Batterham, Hugo J. Bellen

## Abstract

The plight of insect populations around the world and the threats it poses to agriculture and ecosystems has thrown insecticide use into the spotlight. Spinosad is an organic insecticide, considered less harmful to beneficial insects than synthetic insecticides, but its mode of action remains unclear. Using Drosophila, we show that low doses of spinosad reduce cholinergic response in neurons by antagonizing Dα6 nAChRs. Dα6 nAChRs are transported to lysosomes that become enlarged and accumulate upon spinosad treatment. Oxidative stress is initiated in the central nervous system, and spreads to midgut and disturbs lipid storage in metabolic tissues in a Dα6-dependent manner. Spinosad toxicity was ameliorated with the antioxidant N-Acetylcysteine amide (NACA). Chronic exposures lead to mitochondrial defects, severe neurodegeneration and blindness in adult animals. The many deleterious effects of low doses of this insecticide reported here point to an urgent need for rigorous investigation of its impacts on beneficial insects.

## Introduction

The life-cycles of many plant species require pollination by insects, particularly bee species; 75% of crop plants depend on these pollination services to some extent (Klein et al., 2007). Every crop plant species faces the threat of attack by insect pests, typically countered using insecticides targeting proteins that are highly conserved among insect species (Sattelle et al., 2005). While insecticides maximise crop yield, they have the potential to negatively impact populations of insects that provide vital services in agriculture and horticulture (Sánchez-Bayo and Wyckhuys, 2019). There has been a sharp focus on the impact of neonicotinoid insecticides on bees, both in the scientific literature and in public discourse, because of evidence that these chemicals may contribute to the colony collapse phenomenon (Lu et al. 2014; Lundin et al. 2015). Many other insect species are under threat. A recent meta-analysis found an average decline of approximately 9% in terrestrial insect abundance per decade, since 1925 (van Klink et al., 2020), although estimates differ depending on the regions studied and the methodologies used (Wagner et al., 2021). While the extent to which insecticides are involved remains undetermined, they have consistently been associated as a major factor, along with climate change, habitat loss, pathogens and parasites (Cardoso et al., 2020; Sánchez-Bayo and Wyckhuys, 2019; Wagner et al., 2021).

In assessing the risk posed by insecticides, it is important that the molecular and cellular events that unfold following the interaction between the insecticide and its target be understood. Many insecticides target ion channels in the nervous system. At the high doses used to kill pests these insecticides produce massive perturbations to the flux of ions in neurons, resulting in lethality (Perry and Batterham, 2018). But non-pest insects are likely to be exposed to lower doses and the downstream physiological processes that are triggered are poorly understood. In a recent study, low doses of the neonicotinoid imidacloprid were shown to stimulate a constitutive flux of calcium into neurons via the targeted ligand gated ion channels (nicotinic acetylcholine receptors – nAChRs) (Martelli et al., 2020). This causes an elevated level of ROS and oxidative stress which radiates from the brain to other tissues. Mitochondrial damage leads to a significant drop in energy levels, neurodegeneration and blindness (Martelli et al., 2020). Evidence of compromised immune function was also presented, supporting other studies (Chmiel et al., 2019). Many other synthetic insecticides are known to elevate the levels of ROS (Karami-Mohajeri and Abdollahi, 2011; Lukaszewicz-Hussain, 2010; Wang et al., 2016) and may precipitate similar downstream impacts. Given current concerns about synthetic insecticides, a detailed analysis of the molecular and cellular impacts of organic alternatives is warranted. Here we report such an analysis for an insecticide of the spinosyn class, spinosad.

Spinosad is an 85%:15% mixture of spinosyns A and D, natural fermentation products of the soil bacterium *Saccharopolyspora spinosa*. It occupies a small (3%), but growing share of the global insecticide market (Sparks et al. 2017). It is registered for use in more than 80 countries and applied to over 200 crops to control numerous pest insects (Biondi et al., 2012). Recommended dose rates vary greatly depending on the pest and crop, ranging from 96 parts per million (ppm) for Brassica crops to 480 ppm in apple fields (Biondi et al., 2012). Garden sprays containing spinosad as the active ingredient contain doses of up to 5000 ppm. Like other insecticides, the level of spinosad residues found in the field vary greatly depending on the formulation, the application mode and dose used, environmental conditions and proximity to the site of application. If protected from light spinosad shows a half-life of up to 200 days (Cleveland et al., 2002).

Spinosad is a hydrophobic compound belonging to a lipid class known as polyketide macrolactones. Studies using mutants, field-derived resistant strains and heterologous expression have shown that spinosad targets the highly conserved nAChR Dα6 subunit in *Drosophila melanogaster* and a range of other insect species (Perry et al., 2015, 2007; Watson, 2001). This subunit is not targeted by imidacloprid (Watson et al., 2010). The two insecticides differ in their mode of action. Imidacloprid is an agonist causing cation influx into neurons by binding to a site that overlaps with that normally occupied by the native ligand, acetylcholine (ACh) (Buckingham et al., 1997; Martelli et al., 2020; Perry et al., 2008). Spinosad is an allosteric modulator, binding to a site in the C terminal region of the protein (Puinean et al., 2013; Somers et al., 2015). Salgado (1998) measured nerve impulses in cockroaches with electromyograms and found an increased response to spinosad, concluding that spinosad promoted an excitatory motor neuron effect. Salgado and Saar (2004) found that spinosad allosterically activates non-desensitized nAChRs, but that small doses were also capable of antagonizing the desensitized nAChRs. It is currently accepted that spinosad causes an increased sensitivity to ACh in certain nAChRs and an enhanced response at some GABAergic synapses, causing involuntary muscle contractions, paralysis and death (Biondi et al., 2012; Perry et al., 2011; Salgado, 1998). A recent study (Nguyen et al., 2021) showed that both acute and chronic exposures to spinosad causes Dα6 protein levels in the larval brain to decrease. A rapid loss of Dα6 protein during acute exposure was blocked by inhibiting the proteasome system (Nguyen et al., 2021). As *Dα6* loss of function mutants are viable (Perry et al., 2007; Perry et al 2021), it was suggested that the toxicity of spinosad may be due to overloading of protein degradation pathways and/or the internalisation of spinosad where it may cause cellular damage. Spinosad has been shown to cause cellular damage via mitochondrial dysfunction, oxidative stress and programmed cell death in insect cells (*Spodoptera frugiperda* Sf9) (Xu et al., 2018; Yang et al., 2017).

Here we show that while spinosad by itself does not elicit Ca^2+^ flux in *Drosophila* neurons, the response elicited by the cholinergic agonist is stunted upon spinosad pretreatment. Following exposure to spinosad, Dα6 cholinergic receptors traffic to the lysosomes, which induces hallmarks of lysosomal dysfunction. We also show that oxidative stress stemming from lysosomal dysfunction, which is a key factor in spinosad’s mode of action at low doses, triggers a cascade of damage that results in mitochondrial dysfunction, reduced energy levels, extensive neurodegeneration in the central brain and blindness. Given the high degree of conservation of the spinosad target between insect species (Perry et al., 2015), our data suggest that the potential for this insecticide to cause harm in other non-pest insects needs to be thoroughly investigated.

## Results

### Low doses of spinosad affect survival and prevent Ca^2+^ flux into neurons expressing *Dα6* nAChRs

As a starting point to study the systemic effects of low-dose spinosad exposure, a dose that would reduce the movement of third instar larvae by 50% during a 2 hr exposure was determined. This was achieved with a dose of 2.5 ppm (**Figure 1A**). 82% of exposed larvae placed back onto insecticide-free media after being rinsed did not undergo metamorphosis. Death occurred over the course of the next 8 days (**Figure 1B**). Of the 18% of larvae that underwent metamorphosis, only 4% emerged as adults. Pupae showed small and irregular morphology (**Figure 1C**). The effect of this dose was measured on primary culture of neurons expressing the spinosad target, the nAChR Dα6 subunit using the GCaMP5G:tdTomato cytosolic [Ca^2+^] sensor. As no alterations in basal Ca^2+^ levels were detected in response to 2.5 ppm (**Figure 1D, E**), a dose of 25 ppm was tested, again with no measurable impact (**Figure 1D, E**). After 5 min of spinosad exposure, neurons were then stimulated by carbachol, a cholinergic agonist that activates nAChR. Spinosad-exposed neurons exhibited a significant decrease in cholinergic response when compared to non-exposed neurons (**Figure 1D, E**). Total Ca^2+^ content mobilized from ER remained unaltered as measured by thapsigargin-induced Ca^2+^ release (**Figure 1D, E**). These data suggest that spinosad blocks the function of Dα6-containing nAChRs.

**Figure 1.**
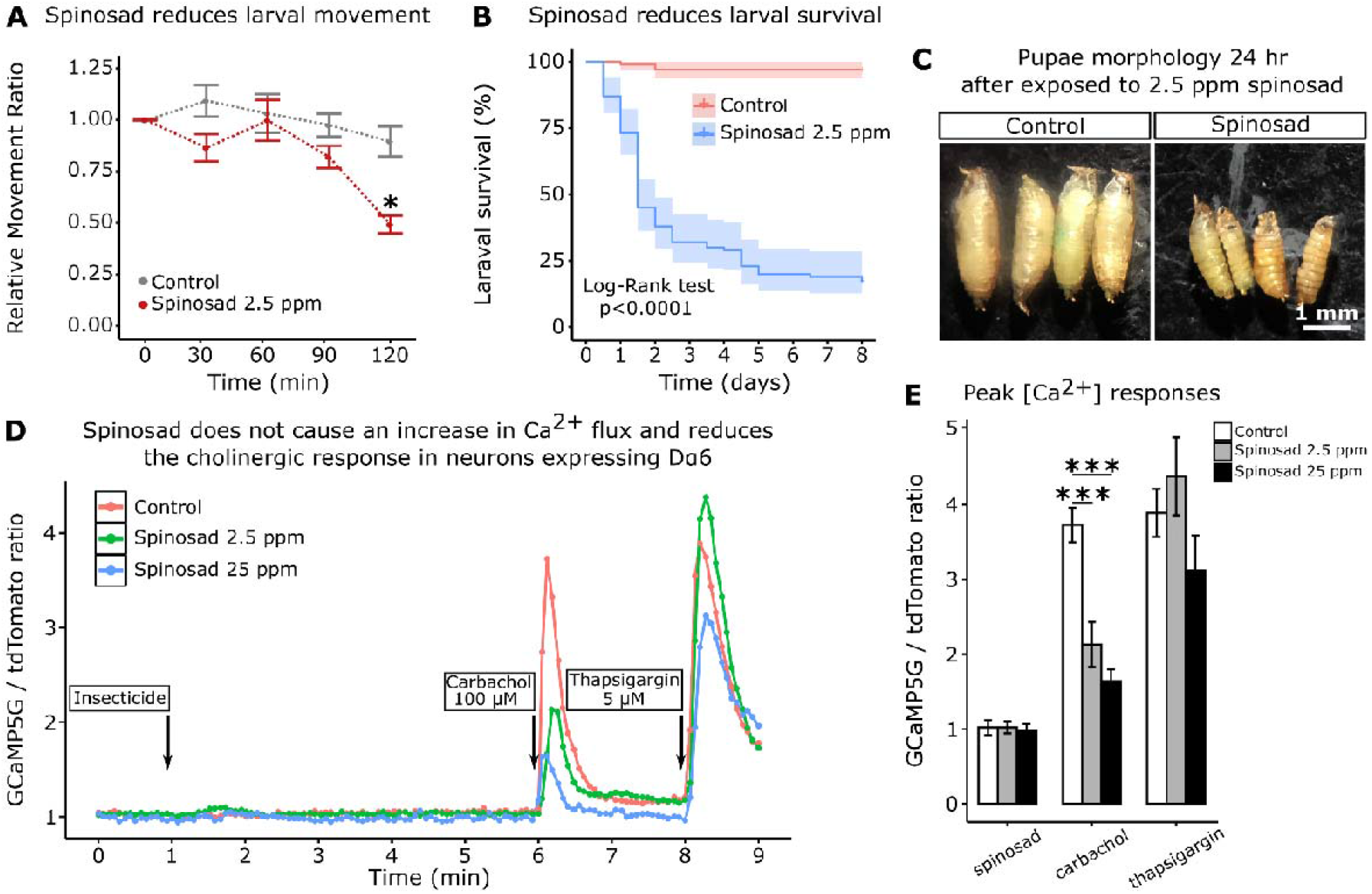
Low doses of spinosad are lethal and fail to increase Ca^2+^ levels in neurons. **A**, Dose response to spinosad by an assay of larval movement over time, expressed in terms of Relative Movement Ratio (RMR); n = 100 larvae/treatment). **B**, % Survival of larvae subjected to a 2 hr exposure to 2.5 ppm spinosad, rinsed and placed back onto insecticide-free medium (n = 100 larvae/treatment). **C**, Pupal morphology, 24 hr after exposure 2.5 ppm spinosad or control solution for 2 hr. **D**, Cytosolic [Ca^2+^] measured by GCaMP in neurons expressing nAChR-Dα6. Measurement is expressed as a ratio of the signals of GCaMP5G signal and tdTomato. Spinosad (2.5 ppm or 25 ppm) was added to the bath solution at 1 minute after recording started. At 6 min and 8 min the spinosad and control groups were stimulated by 100 µM carbachol and 5 µM Thapsigargin, respectively. Each point represents the average of at least 50 cells. **E**, Peak [Ca^2+^] responses to spinosad and carbachol. Error bars represent s.e.m.; shaded areas in **B** represent 95% confidence interval (Kaplan-Meier method and the Log-rank Mantel-Cox test; P < 0.0001). **A** and **E**, t-test; *P < 0.05, ***P <0.001.

### Spinosad exposure causes lysosomal alterations, mitochondrial impairment and increase oxidative stress

To test whether blocked Dα6-containing nAChRs could cause receptor recycling from membrane and thus increase lysosome digestion, LysoTracker staining was used to assess lysosomal function. Whereas no phenotype was observed after 1 hr exposure, a 2 hr exposure to 2.5 ppm spinosad caused an 8-fold increase in the area occupied by lysosomes in the larval brain (**Figure 2A, B**). 6 hr after larvae were subjected to the 2 hr exposure, the area occupied by lysosomes in brains was 24-fold greater than in controls (**Figure 2A, B**). No increase in the area occupied by lysosomes was observed after exposure to imidacloprid, showing that this is a spinosad specific response (**Figure 2 – figure supplement 1**). These observations, in combination with the findings of Nguyen et al. (2021) suggested that binding of spinosad to Dα6 nAChRs may promote their trafficking to lysosomes. To investigate this hypothesis, the brains of larvae expressing a fluorescently (CFP) tagged Dα6 nAChR subunit were stained with LysoTracker. Exposure to 2.5 ppm spinosad showed a significant reduction of the Dα6 CFP signal from neuronal membranes over time and colocalization with lysosomes (**Figure 2C; Figure 2 – figure supplement 2**). Importantly, enlarged lysosomes were not observed in *Dα6 knockout* mutants, regardless of spinosad exposure (**Figure 2 – figure supplement 1**), indicating that the lysosomal expansion is dependent on the presence of Dα6 nAChRs.

**Figure 2.**
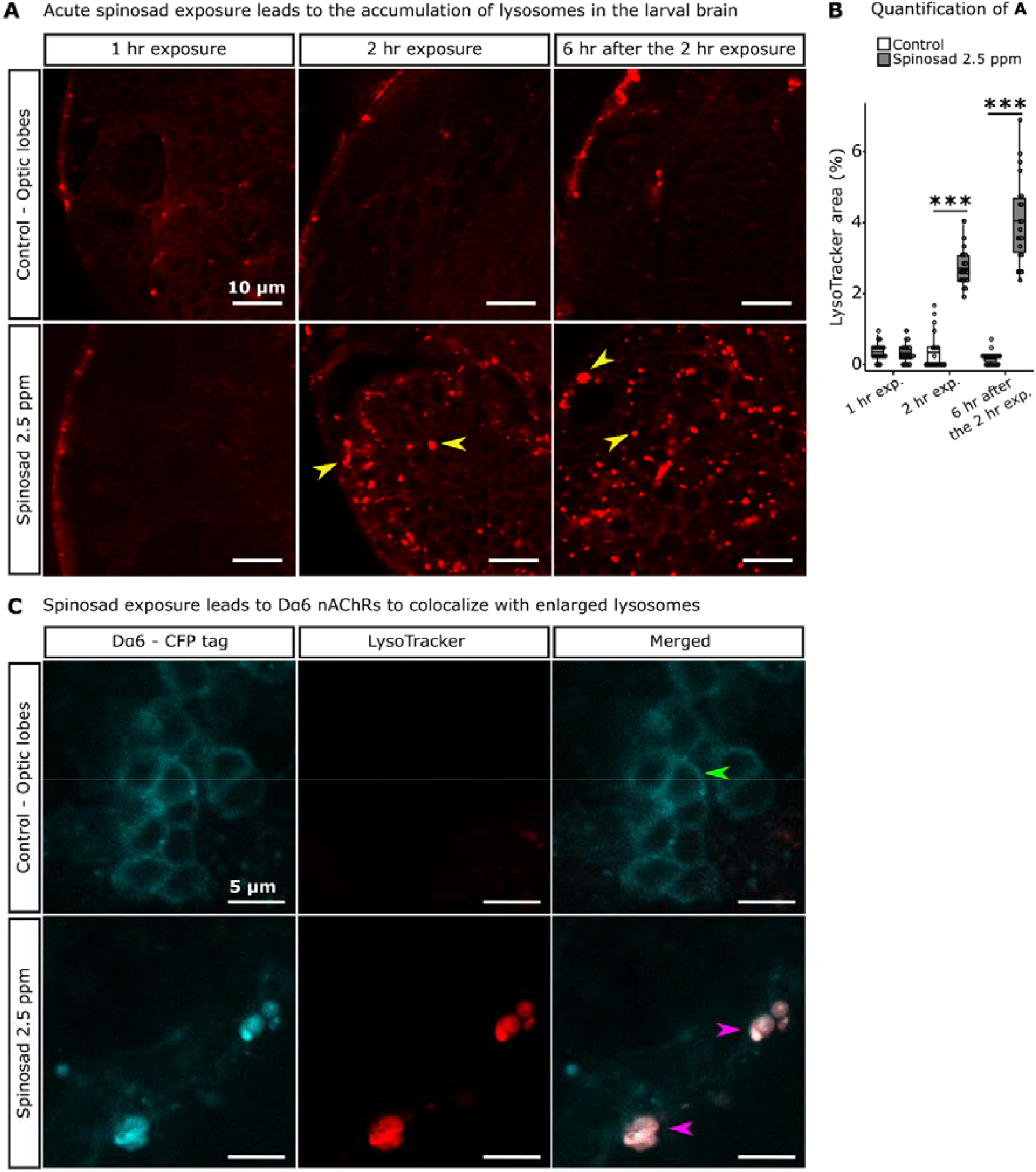
Spinosad exposure causes lysosomal expansion and Dα6 nAChRs colocalize with enlarged lysosomes. **A**, Larvae exposed to 2.5 ppm spinosad for 2hr show a significant increase in the number of enlarged lysosomes in the brain, not observed following a 1hr exposure. 6hrs after the 2hr exposure the number of enlarged lysosomes is further increased. Yellow arrowheads indicate enlarged lysosomes. Lysotracker staining, 400 x magnification. **B**, Lysotracker area in the optic lobes (%) (n = 7 larvae/treatment, 3 optic lobe sections/larva). **C**, Larvae expressing Dα6 tagged with CFP exposed to 2.5 ppm spinosad for 2 hr show co-localization of the Dα6 and lysosomal signals. Green arrowhead indicates Dα6 CFP signal in neuronal membranes of non-exposed larvae. Pink arrowheads indicate Dα6 CFP signal colocalizing with lysosomes. Lysotracker staining, 600 x magnification. Microscopy images obtained in Leica SP5 Laser Scanning Confocal Microscope. t-test; ***P < 0.001.

Defects in lysosomal function have been shown to impact other organelles, especially mitochondria (Deus et al., 2020). To assess mitochondrial dysfunction, we examined the levels of superoxide anion (O_2_^−^), a primary reactive oxygen species (ROS) produced by mitochondria (Valko et al., 2007), using dihydroethidium (DHE) staining. After a 1 hr exposure to 2.5 ppm spinosad, there was a mean 89% increase in O_2_^−^ accumulation in the brain. After 2 hr the levels were lower than at the 1hr time point, but still 44% higher than in the unexposed controls (**Figure 3A, B**). A different pattern was observed in the anterior midgut. A significant increase in accumulation compared with the controls (28%) was only observed at the 2 hr time point (**Figure 3A, B**).

**Figure 3.**
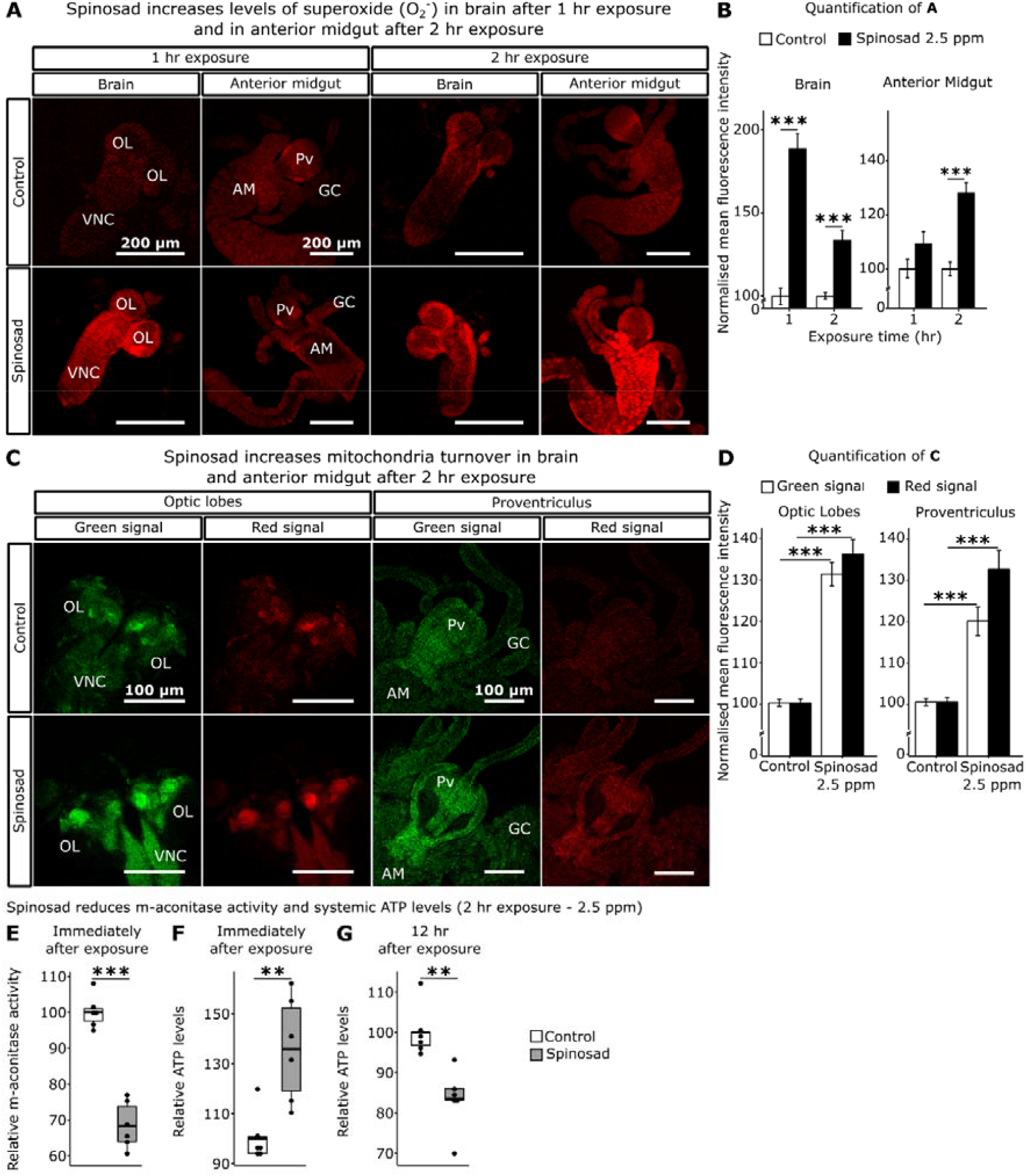
Spinosad exposure impacts ROS levels, mitochondrial turnover and energy levels. **A**, Superoxide levels in the brain and anterior midgut of larvae exposed to 2.5 ppm spinosad for either 1 hr or 2 hr. Tissue stained with DHE. **B**, Normalized mean fluorescence intensity of DHE (n = 15 larvae/treatment; 3 sections/larva). **C**, Optic lobes of the brain and proventriculus of MitoTimer reporter strain larvae. 2.5 ppm spinosad exposure for 2 hr increased the signal of healthy (green) and unhealthy (red) mitochondria (n = 20 larvae/treatment; 3 image sections/larva). **D**, Normalized mean fluorescence intensity of MitoTimer signals. Error bars indicate standard error. **E**, Relative m-aconitase activity in whole larvae (n = 25 larvae/replicate; 6 replicates/treatment) exposed to 2.5 ppm spinosad for 2 hr. **F**, Relative systemic ATP levels immediately after the 2 hr exposure to 2.5 ppm spinosad (n = 20 larvae/ replicate; 6 replicates/ treatment). **G**, Relative systemic ATP levels 12 hr after exposure to 2.5 ppm spinosad (n = 20 larvae/ replicate; 6 replicates/ treatment). OL – optic lobe; VNC – ventral nerve cord; Pv – proventriculus; GC – gastric caeca; AM – anterior midgut. Error bars in **B** and **D** represent mean ± s.e.m. Microscopy images obtained in Leica SP5 Laser Scanning Confocal Microscope, 200x magnification. t-test; **P < 0.01; ***P < 0.001.

Mitochondrial turnover was assessed using the MitoTimer reporter line (Gottlieb and Stotland, 2015). A 2hr spinosad exposure induced an increase of 31% and 36% for the green (healthy mitochondria) and red (stressed mitochondria) signals in the optic lobes of the larval brain, respectively (**Figure 3C, D**). For the digestive tract, a 19% and 32% increase were observed in the proventriculus for green and red signal, respectively (**Figure 3C, D**). To examine the impact of ROS we measured the enzyme activity of mitochondrial aconitase, a highly ROS sensitive enzyme (Yan et al., 1997). We observed a mean 34% reduction in aconitase activity (**Figure 3E**), indicating an increased presence of ROS in mitochondria during the 2 hr exposure. Immediately after the 2 hr exposure, a mean 36% increase in systemic ATP levels was observed (**Figure 3F**), followed by a 16.5% reduction 12 hr after the 2 hr exposure (**Figure 3G**). The initial increase in energy levels is consistent with the increase in the green signal observed with MitoTimer at this time point. However, the reduction in ATP levels 12 hr after the exposure shows that the mitochondrial energy output is eventually impaired.

### Oxidative stress created by spinosad affects lipids, motility, and survival

Oxidative stress has the ability to affect the lipid environment of metabolic tissues, causing bulk redistribution of lipids into lipid droplets (LD) (Bailey et al., 2015). An elevation of ROS levels in the *Drosophila* larval brain has been shown to cause an increase in LD numbers in the fat body as well as a decreases LD in the midgut and Malpighian tubules (Martelli et al., 2020). The impact of spinosad on LD numbers was therefore examined. Larvae exposed to 2.5 ppm spinosad for 2 hr showed a 52% increase in the area covered by LD in the fat body (**Figure 4A, B**), with a significant reduction in the number of large LD and an increase in small LD (**Figure 4 – figure supplement 1**). Pre-treatment with the antioxidant N-Acetylcysteine amide (NACA) significantly reduced the impacts of spinosad exposure on this phenotype. Even though still significant, the area occupied by LD in fat bodies increased only 20% with NACA pre-treatment (**Figure 4A, B**). Antioxidant pre-treatment also significantly improved movement of larvae exposed to spinosad (**Figure 4C**), and survival, which increased from 4% to 15% (**Figure 4D**).

**Figure 4.**
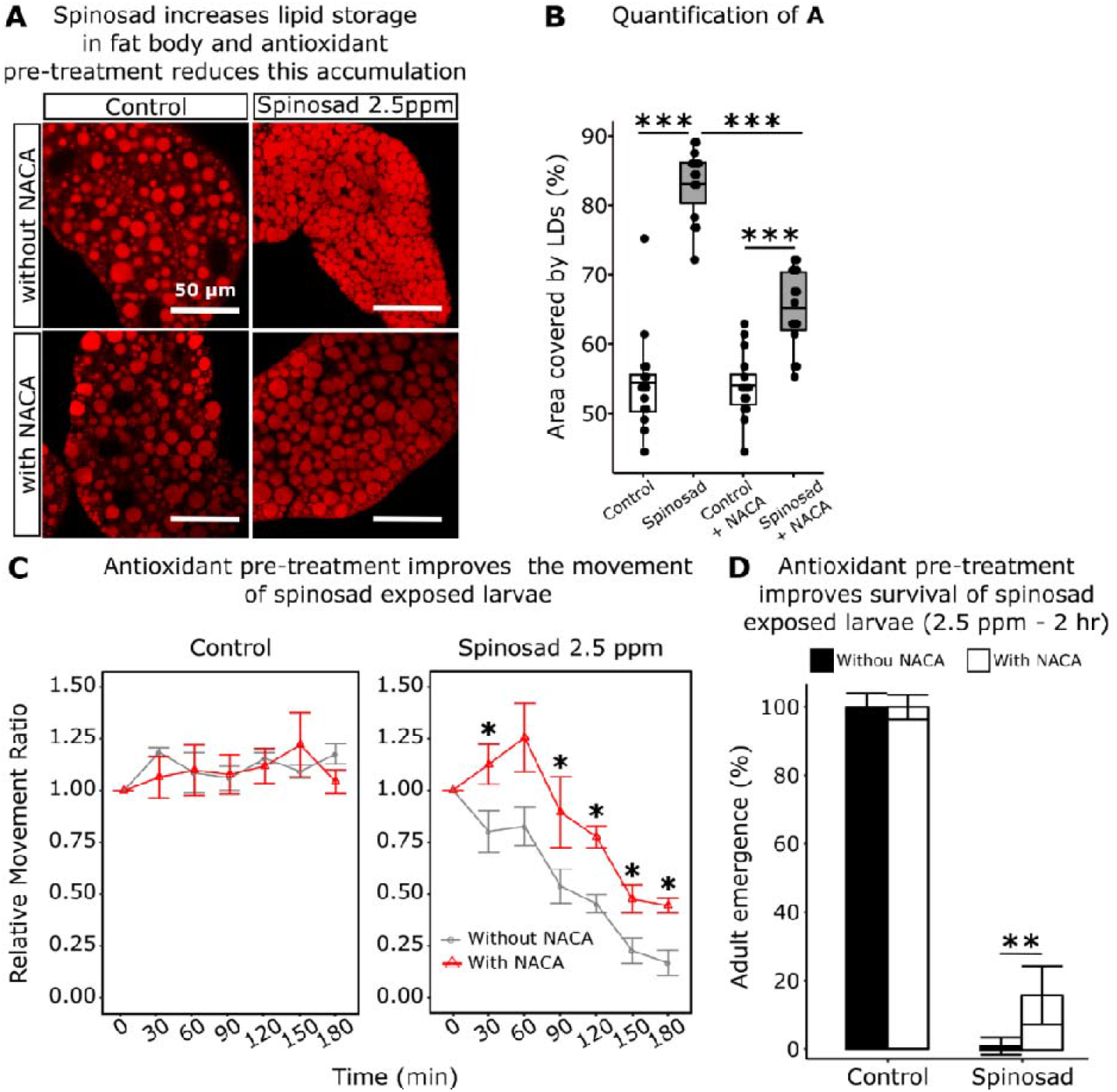
Spinosad increases lipid storage in fat body. Antioxidant pre-treatment reduces this accumulation and improves larval movement and survival. **A**, Larvae exposed to 2.5 ppm spinosad for 2 hr show an accumulation of LD in the fat body. A 5 hr pre-treatment with 300 µg/mL of antioxidant N-acetylcysteine amide (NACA) reduces this accumulation. Nile red staining. Images obtained using a Leica SP5 Laser Scanning Confocal Microscope, 400x magnification. **B**, Percentage of area occupied by LD in fat body (n = 3 larvae/treatment; 5 image sections/larva). **C**, Pre-treatment with NACA improves the movement of spinosad exposed larvae. Dose response to insecticide analysed using the Wiggle Index analysis. Results are expressed in terms of Relative Movement Ratio (RMR) values as a function of exposure time in minutes (n = 25 larvae/replicate; 4 replicates/ treatment). **D**, Pre-treatment with NACA improves survival of larvae exposed to spinosad. Corrected adult emergence (%) (n = 100 larvae/ treatment). Bars indicate corrected percentage survival (Abbots’ correction). Error bars in **C** represent the s.e.m. and in **D** the 95% confidence interval. t-test; *P < 0.05; **P < 0.01; ***P < 0.001.

In order to test whether doses that do not impact survival could also cause similar perturbations to the lipid environment, sublethal acute doses were determined. Larvae exposed to 0.5 ppm for 2 hr or 0.1 ppm for 4 hr showed no impact in adult eclosion after being rinsed and placed back onto insecticide-free media (**Figure 4 – figure supplement 2**). Both doses caused on average a 29% increase in the area occupied by LD in fat bodies (**Figure 4 – figure supplement 2**). That this impact is smaller than that observed for the 2.5 ppm shows that this phenotype is dose dependent. Once again, an increase in the number of small LD and reduction in the number of large LD was observed (**Figure 4 – figure supplement 3**). Using these sublethal doses, other metabolic tissues were investigated. The doses of 0.5 ppm for 2 hr and 0.1 ppm for 4 hr caused a mean 72% and 73% reduction in the total number of LD in the Malpighian tubules, respectively (**Figure 4 – figure supplement 2**). We also identified a reduction in the numbers of LD in the LD region of the posterior midgut (**Figure 4 – figure supplement 4**).

### A brain signal triggers the impacts of spinosad on metabolic tissues

Once inside the insect body, spinosad could theoretically access any tissue via the open circulatory system. Given that the target Dα6 nAChRs are localized in the brain (Perry et al., 2015; Somers et al., 2015), and that elevated levels of ROS were observed earlier in the brain than in metabolic tissues, prompts a significant question. Could the interaction between spinosad and Dα6 in the brain provide the signal that ultimately leads to the observed disturbance of the lipid environment in the metabolic tissues? Two different *Dα6 knockout* mutants (Line 14 *Dα6 KO* and Canton S *Dα6 KO*) and their respective genetic background control lines (Line 14 – used in experiments so far, and Canton S) were tested. Larvae were exposed to 2.5 ppm of spinosad for 2 hr. Neither of the mutants tested showed an increase in the area occupied by LD, compared to their respective background lines, under conditions of spinosad exposure (**Figure 5A-D**). We also quantified the level of lipids in hemolymph. Whereas Line 14 and Canton S showed an average 10% and 13% increase in response to spinosad, respectively, neither of the *Dα6 KO* mutants showed significant changes (**Figure 5E**). Hence, *Dα6* mediates the observed lipid phenotypes.

**Figure 5.**
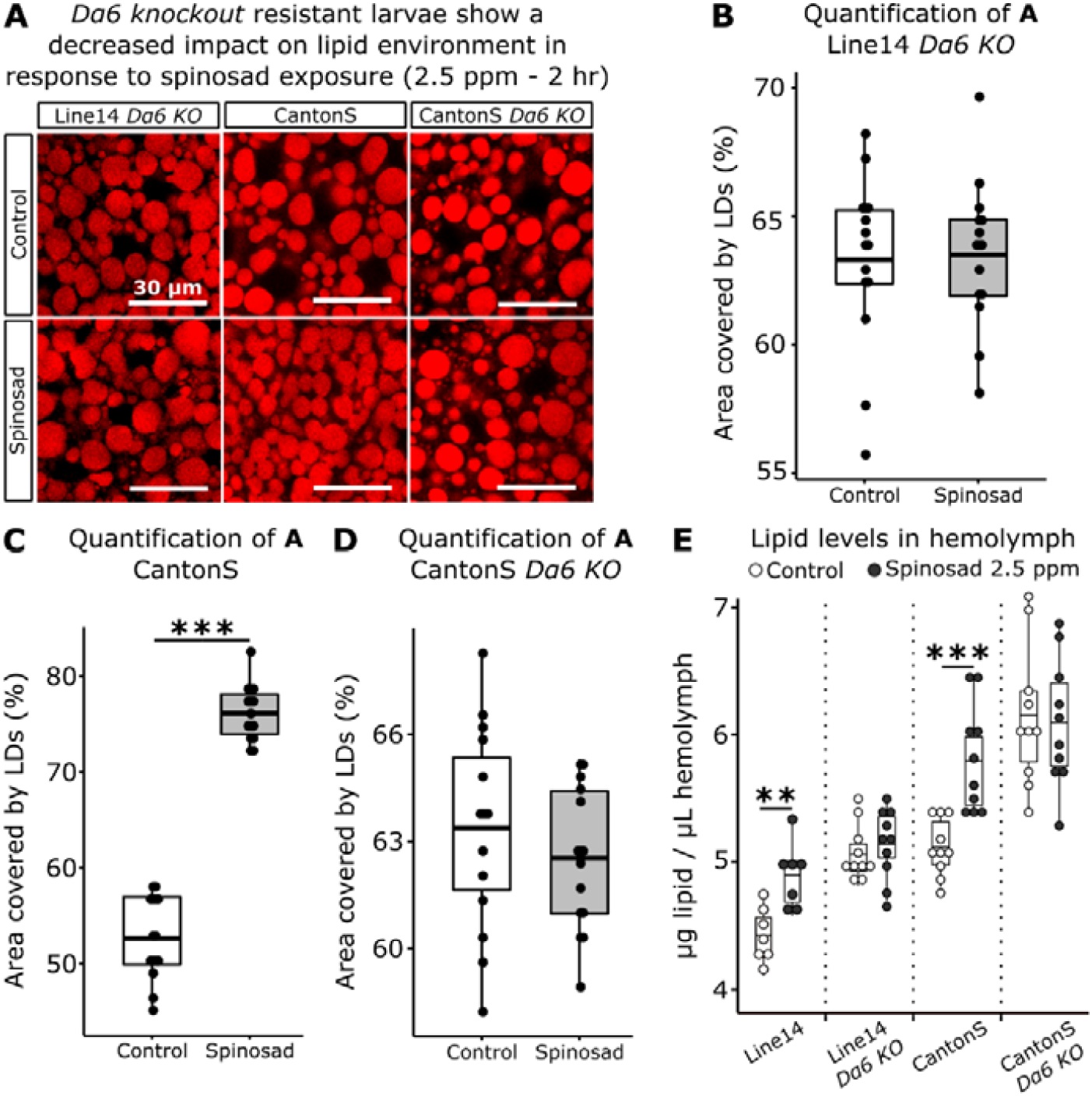
*Dα6 knockout* (*KO*) resistant larvae show a decreased impact on lipid environment in response to spinosad exposures. **A**, Larvae exposed to 2.5 ppm of spinosad for 2 hr. Nile red staining. Images obtained in Leica SP5 Laser Scanning Confocal Microscope, 400x magnification (n = 3 larvae/ treatment; 5 image sections/larva). **B**, Percentage of area occupied by LD in fat body of Line 14 *Dα6 KO* larvae. **C**, Percentage of area occupied by LD in fat body of Canton S larvae. **D**, Percentage of area occupied by LD in fat body of Canton S *Dα6 KO* larvae. **E**, Amount of lipids in hemolymph (µg/µL) of Line 14 *Dα6 KO*, Canton S and Canton S *Dα6 KO* larvae exposed to 2.5 ppm spinosad for 2 hr. Measured using the colorimetric vanillin assay (n = 10 replicates/treatment/time-point; 30 larvae/replicate). t-test; ***P < 0.001.

### Spinosad triggers major alterations in the lipidome pointing to impaired membrane function and decreased mitochondrial cardiolipins

To further investigate the impacts on the lipid environment we performed a lipidomic analysis on whole larvae exposed to 2.5 ppm spinosad for 2 hr. Significant changes were observed in the levels of 88 lipids out of the 378 detected by mass spectrometry (**Figure 6A**; **Figure 6 – table supplement 1**). A significant portion of the changes in lipids correspond to a reduction in phosphatidylcholine (PC), phosphatidylethanolamine (PE) and some triacylglycerol (TAG) species. Multivariate analysis (**Figure 6B**) indicates that the overall lipidomic profiles of exposed larvae forms a tight cluster that is distinct from the undosed control. The use of whole larvae for lipidomic analysis reduces the capacity to detect significant shifts in lipid levels that predominantly occur in individual tissues but allows the identification of broader impacts on larval biology. In this context, the observed 65% reduction in the levels of identified cardiolipins (CL) is particularly noteworthy (**Figure 6C**). CL are mostly present in mitochondria and are required for the proper function of the TCA cycle proteins, especially those of Complex 1, the major ROS generator when dysfunctional (Quintana et al., 2010; Ren et al., 2014).

**Figure 6.**
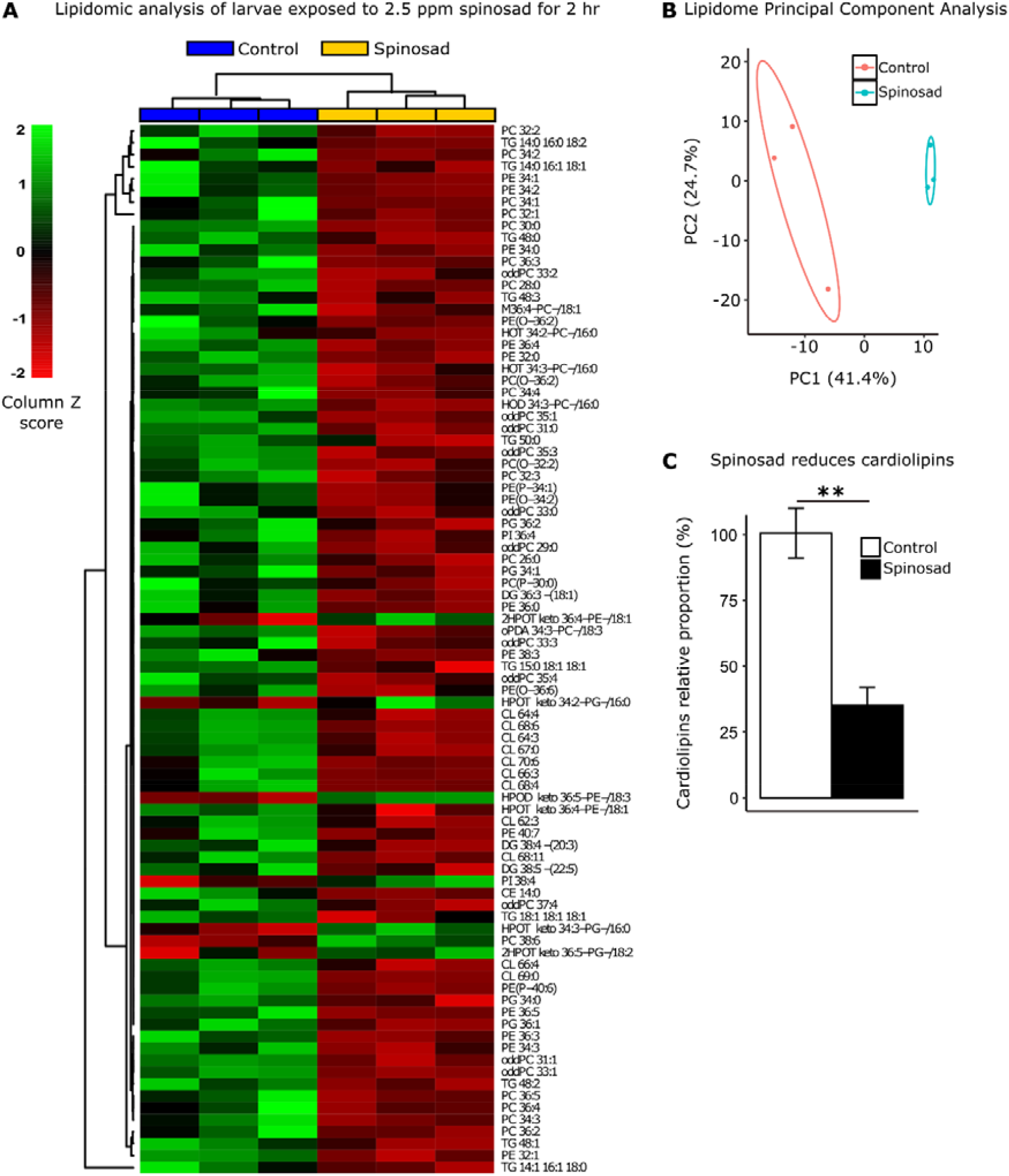
Spinosad disturbs the lipid profile of exposed larvae. Lipidomic profile of larvae exposed to 2.5 ppm spinosad for 2 hr (n = 10 larvae/replicate; 3 replicates/treatment). **A**, 88 lipid species out of the 378 identified were significantly affected by insecticide treatment (One-way ANOVA, Turkey’s HSD, P < 0.05). The column Z score is calculated subtracting from each value within a row the mean of the row and then dividing the resulting values by the standard deviation of the row. The features are color coded by row with red indicating low intensity and green indicating high intensity. **B**, Principal Component Analysis of 378 lipid species. Each dot represents the lipidome data sum of each sample. First component explains 41.4% of variance and second component explains 24.7% of variance. **C**, Relative proportion of cardiolipins in exposed animals versus control. Error bars in **C** represent mean ± s.e.m. t-test; **P < 0.01.

### Chronic low exposure to spinosad causes neurodegeneration and progressive loss of vision

Next, we investigated the effects of chronic exposure to spinosad in adults. A dose of 0.2 ppm spinosad kills 50% of adult female flies within 25 days (**Figure 7A**). Two different behavioural assays were initially assessed: bang sensitivity and climbing. Exposure to 0.2 ppm spinosad for 10 and 20 days increased the bang sensitivity phenotype that has been associated with perturbations in synaptic transmission (Saras and Tanouye, 2016) that can arise from various defects including defective channel localization, neuronal wiring and mitochondrial metabolism (Fergestad et al., 2006) (**Figure 7B**). This assay measures the time it takes for flies to recover to a standing position following mechanical shock induced by vortexing the flies. Exposed flies also performed poorly in climbing assays, a phenotype which is often linked to neurodegeneration (McGurk et al., 2015). Indeed, 16%, 73% and 84% of flies failed to climb after 1, 10 and 20 days of exposure, respectively (**Figure 7C**). These data suggest that low doses of spinosad induce neurodegenerative phenotypes.

**Figure 7.**
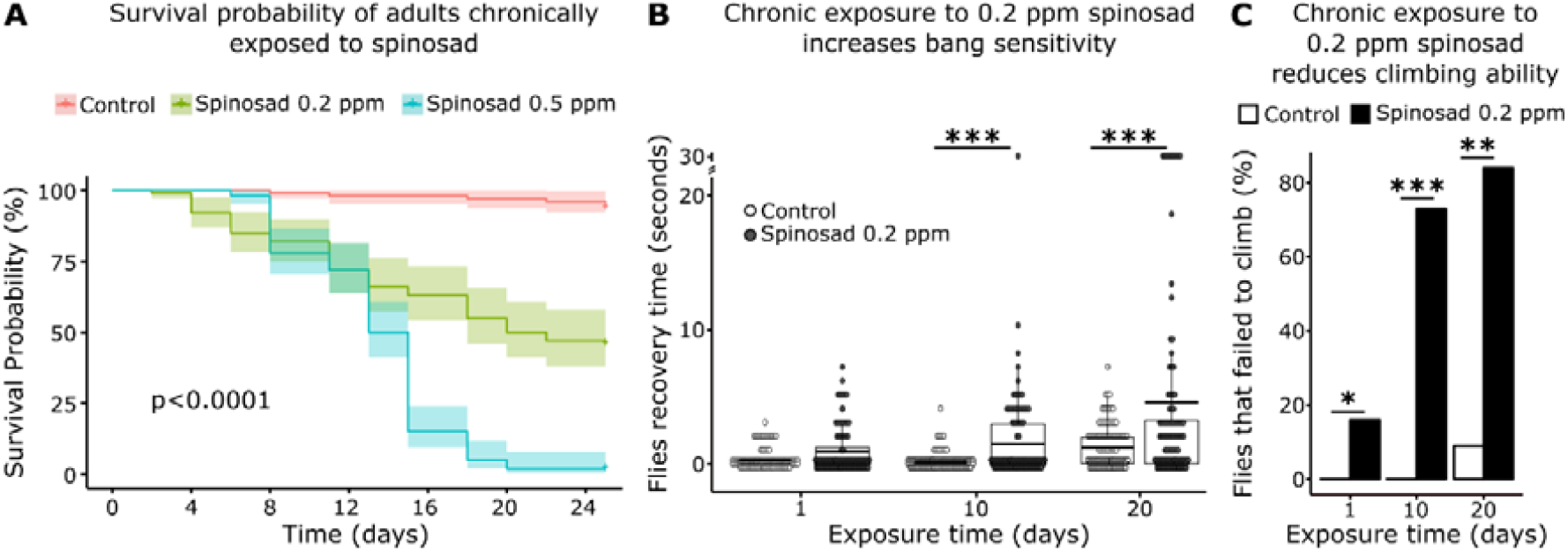
Chronic exposure to spinosad affects behavior. **A**, Determination of a chronic exposure dose that kills 50% of adults within 25 days. Females adults (2-5 days old) were exposed to different concentrations of spinosad for 25 days (n = 25 flies/ replicate; 4 replicates/ treatment). The dose of 0.2 ppm was selected for assessing the impacts of adult chronic exposures. Shaded areas represent 95% confidence interval (Kaplan-Meier method and the Log-rank Mantel-Cox test). **B**, Chronic exposure to 0.2 ppm spinosad increases bang sensitivity. Bang sensitivity assay of adults after 1, 10 and 20 days of exposure. Groups of 5 flies were vortexed in a clear vial for 10 seconds at maximum speed and the recovery time (time regain normal standing posture) for each fly was recorded (n = 100 flies/time point/ treatment). **C**, Chronic exposure to 0.2 ppm spinosad reduces climbing ability. Percentage of adult flies that failed to climb after 1, 10 and 20 days of exposure (n = 100 flies/ time point/ treatment). **B** and **C**, Wilcoxon test; **P < 0.01; ***P < 0.001.

The retina of adult female flies chronically exposed to 0.2 ppm spinosad were examined for evidence of neurodegeneration, such as the accumulation of LD in glial cells based on Nile Red staining (Liu et al., 2015). Nile Red positive accumulations, likely to represent small LD, were observed decorating the plasma membrane of photoreceptor cells (**Figure 8A, B**). Even though nAChR *Dα6* is not expressed in the retina, it is widely expressed in the adult brain, including the lamina, tissue adjacent to the retina where the photoreceptors synapse (**Figure 8 – figure supplement 1**). Indeed, several laminar neurons synpase with the photoreceptors. The accumulation of LD in neurons suggest that the postsynaptic cells that express D6 somehow affect lipid production in PR.

**Figure 8.**
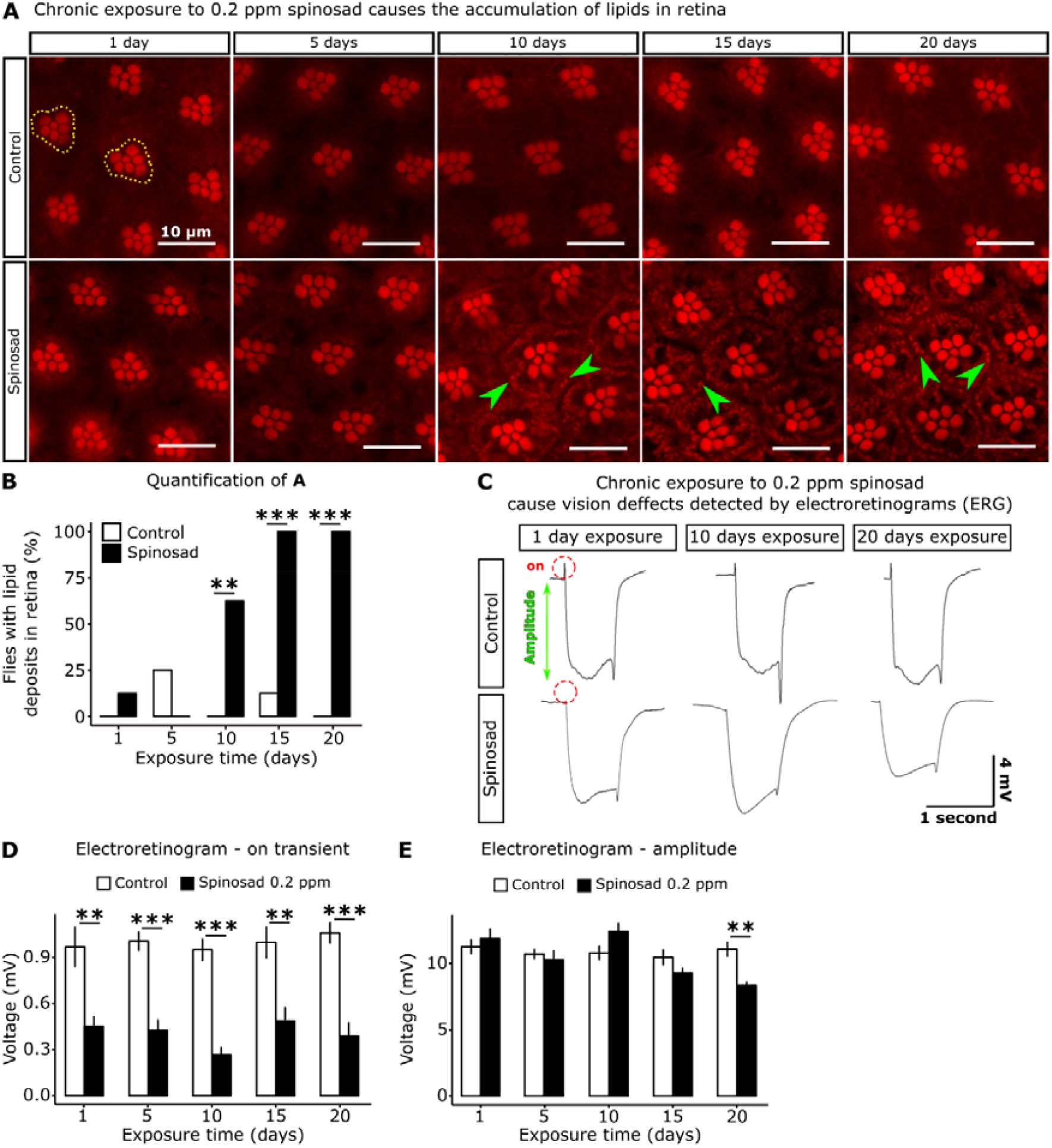
Chronic exposure to spinosad causes loss of vision. **A**, Clusters of rhabdomeres in the retina. In day 1 – control, two clusters of rhabdomeres are delimited with yellow dotted-lines. A diffuse lipid accumulation is observed from day 10 onwards. Nile red staining. 600 x magnification. **B**, Percentage of animals that shows lipid deposits in the retina (n = 8 flies/treatment/time point). **C**, Electroretinograms (ERGs) of animals exposed to 0.2 ppm spinosad for 1, 10 and 20 days. Red dotted circles indicate the on-transient signal and green arrow indicates the amplitude, (n = 8 to 10 adult flies/time point/treatment). **D**, On-transient signal of ERGs after days 1, 5, 10, 15 and 20 of exposure to 0.2 ppm spinosad. **E**, Amplitude of ERGs after days 1, 5, 10, 15 and 20 of exposure to 0.2 ppm spinosad. Microscopy images obtained in Leica SP5 Laser Scanning Confocal Microscope. t-test; **P < 0.01; ***P < 0.001.

To quantify possible impacts on visual function, electroretinograms (ERGs) were performed at regular intervals over the 20 days of exposure (**Figure 8C-E**). ERG recordings measure impulses induced by light. The on-transient is indicative of synaptic transmission between photoreceptor neurons (PR) and postsynaptic cells, whilst the amplitude measures the phototransduction cascade (Wang and Montell, 2007). A large reduction in the on-transient was observed from day 1 of exposure, whereas the amplitude was only significantly impacted after 20 days of exposure. The reduction in the on-transient is evidence of a rapid loss of synaptic transmission in laminar neurons (Wang and Montell, 2007) and hence impaired vision after just one day of exposure.

To investigate the ultrastructure of the PR synapses we used Transmission Electron Microscopy. Severe morphological alterations were detected in transverse sections of the lamina of flies exposed for 20 days (**Figure 9A-F**). Vacuoles of photoreceptor terminals or postsynaptic terminals of synapsing neurons were observed in the lamina cartridges (**Figure 9B**). On average 70% of images showed the presence of vacuoles in lamina cartridges (**Figure 9E**). Large intracellular compartments were also observed in the dendrites of the postsynaptic neurons in the lamina (**Figure 9B-D**). These do not correspond to normal structures found in healthy lamina (**Figure 9A**). The lamina of exposed flies also showed a mean 34% increase in the number of mitochondria (**Figure 9F**), many of which appear defective (**Figure 9B**). In examining the visual system of a *Dα6 KO* mutants reared without spinosad, mild impacts were identified in ERG amplitude but a very significant reduction in on-transient was observed, consistent with a requirement for *Dα6* in postsynaptic cells of the photoreceptors. No morphological alterations were detected in the lamina by TEM (**Figure 9 – figure supplement 1**).

**Figure 9.**
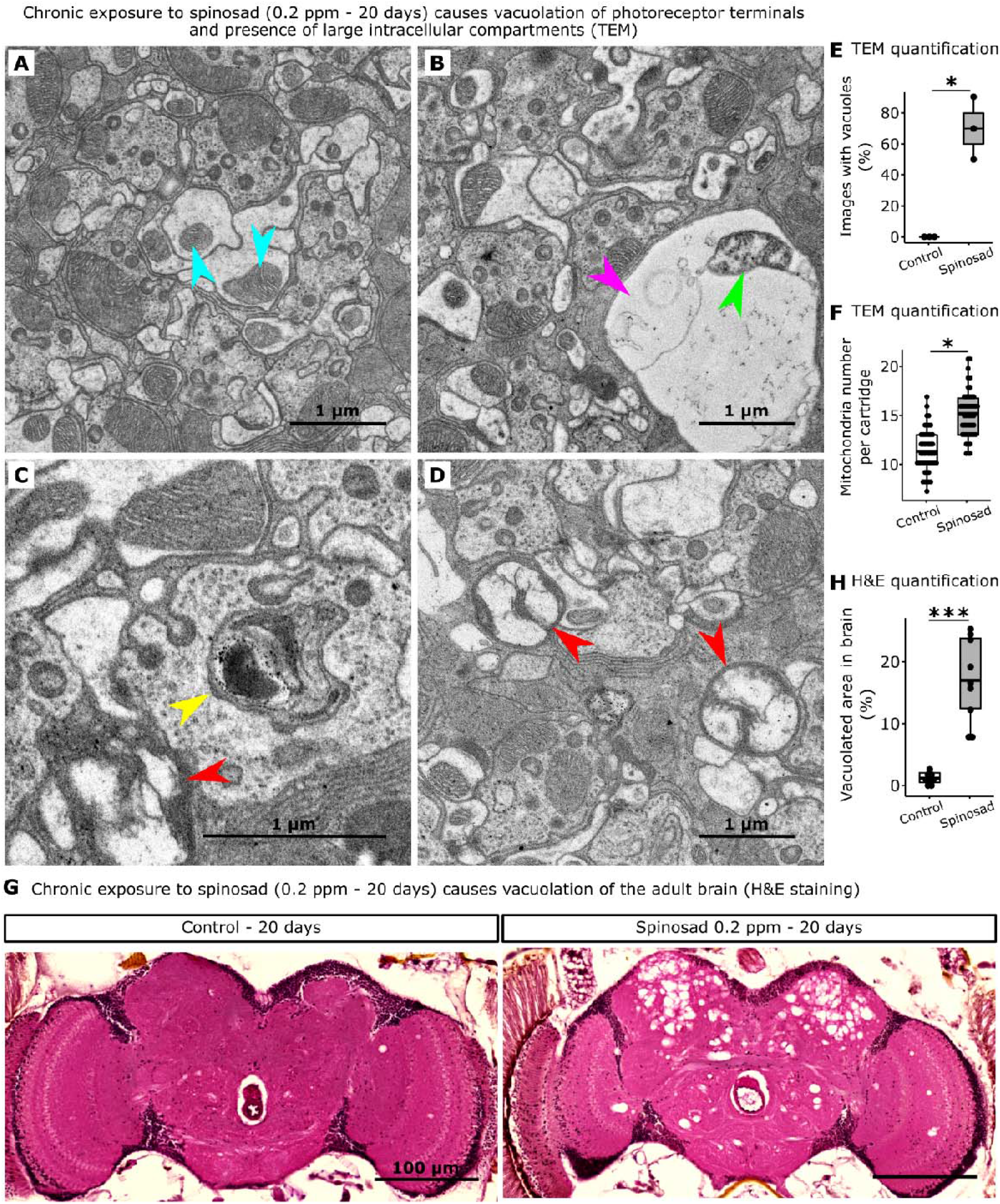
Chronic exposure to spinosad causes neurodegeneration. **A**, Transmission electron microscopy (TEM) of the lamina of a control animal showing a regular cartridge, blue arrowheads indicate normal mitochondria. **B-D**, TEM of lamina of flies exposed to 0.2 ppm spinosad for 20 days. **B**, Pink arrowhead indicates vacuole and green arrowhead indicates a defective mitochondrion. **C**, Yellow arrowhead indicates an enlarged digestive vacuole inside a photoreceptor terminal. **D**, Red arrowheads indicate the presence of large unidentified intracellular compartments. **E**, Percentage of images showing vacuoles in lamina cartridges (10 images/fly; 3 flies/ treatment). **F**, Number of mitochondria per cartridge (n = 3 flies/group; 16 cartridges/fly). **G**, Flies exposed to 0.2 ppm spinosad for 20 days show vacuolation of the central brain. Brain frontal sections stained with hematoxylin and eosin (H&E). **H**, Quantification of neurodegeneration in terms of percentage of brain area vacuolated (n = 3 flies/treatment). t-test; *P < 0.05; ***P < 0.001.

Lastly, Hematoxylin & Eosin stain (H&E) of adult flies painted a picture of the neurodegeneration caused outside the visual system by chronic low dose exposure to spinosad. 20 days of exposure caused numerous vacuoles in the central brain (**Figure 9G, H**). On average, 17% of the total central brain area was consumed by vacuoles in exposed flies.

## Discussion

### Spinosad antagonizes neuronal activity

In this study we provide evidence of the mechanism and consequences of exposure to low doses of spinosad. This organic insecticide leads to a lysosomal dysfunction associated with a mitochondrial dysfunction, elevated levels of ROS, lipid mobilization defects and neurodegeneration. Spinosad has been characterized as an allosteric modulator of the activity of its primary target, the nAChR – Dα6 subunit, causing fast neuron over-excitation (Salgado, 1998). Here, the capacity of spinosad to interact with its target nAChRs to stimulate the flux of Ca^2+^ into neurons was quantified. The results obtained with the GCaMP assay showed that spinosad caused no detectable increase or decrease in Ca^2+^ flux into *Dα6* expressing neurons, but it reduced the cholinergic response (**Figure 1**). Given that spinosad binds to the C terminal region of the protein (Crouse et al., 2018; Puinean et al., 2013; Somers et al., 2015), these findings are consistent with a non-competitive antagonist mode of action for spinosad on nAChRs. That *Dα6* loss of function mutants are viable (Perry et al., 2007) creates a conundrum that can be resolved if a significant component of spinosad’s toxicity is due to molecular events that play out elsewhere in the cell. Blocked neuronal receptors can be recycled from the plasma membrane through endocytosis (Saheki and De Camilli, 2012). Our data indicate that spinosad exposure leads to the removal of Dα6 nAChRs from neuronal membranes (Nguyen et al., 2021) and localization to enlarged lysosomes, resulting in lysosomal expansion (**Figure 2C**) and lysosomal dysfunction.

### Spinosad causes lysosomal storage diseases - like phenotype

The following observations suggest that spinosad induces lysosomal dysfunction. LysoTracker staining reveals a very significant accumulation of enlarged lysosomes in the brain in response to spinosad, but not in the presence of imidacloprid, another insecticide which also binds to nAChRs (**Figure 2 – figure supplement 1**). Importantly, *Dα6 knockout* flies show no accumulation of LysoTracker staining, clearly showing that the lysosomal lesions rely on the presence of Dα6 and spinosad (**Figure 2 – figure supplement 1**). Whether spinosad molecules are ferried to lysosomes along with Dα6 subunits and accumulate into these organelles remains to be clarified. However, the increased severity in the lysosomal phenotype after exposure ceases (**Figure 2A, B**) is consistent with the poisoning of these organelles. Lysosomes become enlarged as they accumulate undigested material, which can lead to recycling problems for neurons (Darios and Stevanin, 2020). If spinosad remains bound to the receptor and is ferried into the lysosomes it may contribute to a lysosomal dysfunction akin to Lysosomal Storage Disease (LSD) (Darios and Stevanin, 2020). To date there is little published evidence of spinosad metabolites in insects. Spinosad is a complex polyketide macrolactone that may not be hydrolysed by lysosome acidic enzymes and could accumulate in the lumen of these organelles.

Our hypothesis for the mode of action of spinosad is illustrated in **Figure 10**. Spinosad exposure shows a delayed effect on larval movement when compared to imidacloprid (Denecke et al., 2015; Martelli et al., 2020). We attribute this to the time taken for a threshold level of lysosomal damage to accumulate. Imidacloprid is readily metabolized and the metabolites are excreted (Fusetto et al., 2017), leaving little lingering damage. In contrast, following a 2hr exposure to 2.5ppm spinosad, 3^rd^ instar larvae show a developmental arrest and die after several days (**Figure 1**). The LSD-like dysfunction is also likely the underlying cause for the severe vacuolation of adult central brain under spinosad chronic exposure. Recycling defects in neuronal cells caused by LSD impair cell function, ultimately triggering neurodegeneration (Darios and Stevanin, 2020). Nguyen et al. (2021) recently showed that flies treated with a proteasome inhibitor drug, bortezomib, present with a reduced loss of Dα6 from neuronal membranes when exposed to spinosad. That suggests that the proteasome degradation pathway could also be involved in recycling spinosad-blocked Dα6 subunits. Receptors marked for proteasome degradation can end up in lysosomes as these pathways engage in crosstalk (Korolchuk et al., 2010).

**Figure 10.**
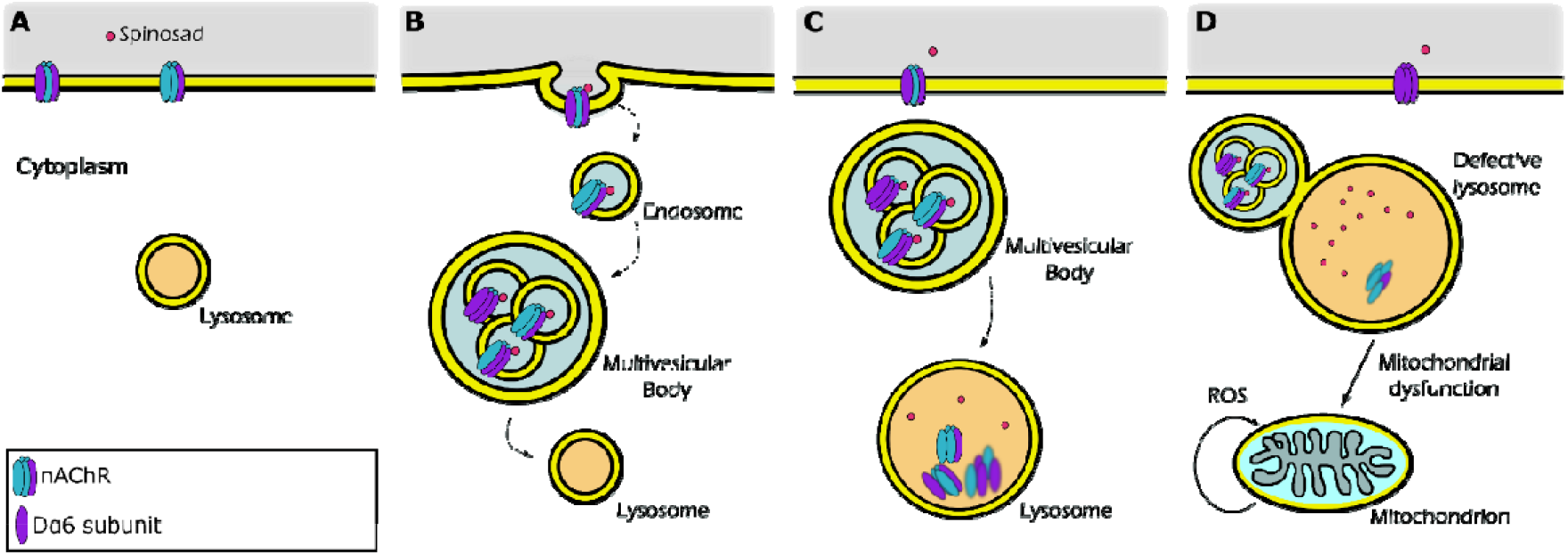
Proposed mechanism for internalization of spinosad after binding to the Dα6 nAChR target. **A**, Spinosad binds to Dα6 subunit of nAChRs in the neuronal cell membranes. **B**, The binding of spinosad leads to Dα6 nAChR blockage, endocytosis and trafficking to lysosome. **C**, Spinosad accumulates in lysosomes, while receptors and other membrane components are digested. **D**, Enlarged lysosomes due to accumulation of undigested material do not function properly leading to cellular defects which may include mitochondrial dysfunction, increased mitochondrial ROS production and eventually cell vacuolation and neurodegeneration.

### Spinosad triggers oxidative stress

Extensive evidence connects lysosomal disorders with mitochondrial dysfunction (Plotegher and Duchen, 2017; Stepien et al., 2020; Yambire et al., 2018). Mitochondrial dysfunction is widespread in LSD and is involved in its pathophysiology. Although mitochondrial dysfunction in LSD seems to have a multifactorial origin, the exact mechanisms remain unclear. Lysosomal disorders may lead to cytoplasmic accumulation of toxic macromolecules, impaired degradation of damaged mitochondria and dysregulation of intracellular Ca^2+^ homeostasis, resulting in increased ROS generation and reduced ATP levels (Plotegher and Duchen, 2017). The severe lysosomal dysfunction observed here is the most likely cause for the mitochondrial defects and increased ROS generation triggered by spinosad exposure.

The evidence for oxidative stress produced during spinosad exposure comes from the accumulation of superoxide, increased mitochondrial turnover, reduced activity of the ROS sensitive enzyme m-aconitase and reduced ATP levels (**Figure 3**), accumulation of LD in fat bodies (**Figure 4**), and severe reduction of cardiolipin levels that typically associated with defects in the electron transport chain and increased ROS production (Quintana et al., 2010) (**Figure 6**). Increasing levels of ROS in the larval brain using RNAi has been shown to disturb mitochondrial function triggering changes in lipid stores in metabolic tissues (Martelli et al., 2020). Oxidative stress promotes redistribution of membrane lipids into LD, reducing their susceptibility to lipid peroxidation (Bailey et al., 2015). Here, increases in lipid stores were observed in the fat body, with a reduction in the numbers of large LD and accumulation of small LD, a reduction in LD in the Malpighian tubules and midgut and changes in lipid levels in the hemolymph (Figure (**Figure 4**). Our lipidome analysis revealed reduction of PE and PC levels (**Figure 6**), consistent with impaired membrane fluidity and altered LD dynamics (Dawaliby et al., 2016; Guan et al., 2013; Krahmer et al., 2011).

The use of the antioxidant NACA reduces the accumulation of LD in the fat body linking this phenotype to oxidative stress (**Figure 4**). NACA also diminished spinosad toxicity by reducing the impact on larval movement and survival (**Figure 4**). *Dα6 knockout* mutants exposed to spinosad show no accumulation of LD in the fat body or change of lipid levels in hemolymph indicating that these phenotypes are due to the spinosad:Dα6 interaction (**Figure 5**). Exposed to 7.7 ppb (parts per billion) for 24 hr was shown to cause the vacuolation of epithelial cells of the midgut and Malpighian tubules of honeybees (*Apis mellifera*) (Lopes et al., 2018). It is not clear whether this is due to the spinosad:Dα6 interaction precipitating elevated levels of ROS.

A striking similarity between impacts caused in metabolic tissues by spinosad and imidacloprid (Martelli et al., 2020) is observed, although the impacts induced by spinosad are more severe. In the case of imidacloprid, these perturbations were shown to be caused by an oxidative stress signal initiated by an increase Ca^2+^ influx into neurons caused by the insecticide binding to its nAChR targets (Martelli et al., 2020). It was proposed that peroxidised lipids generated in the brain and carried in hemolymph precipitate oxidative damage to other tissues (Ioannou et al., 2019; Valko et al., 2007). Concomitantly, *Da6* has been associated with the response to oxidative stress. *Da6* mutants are more susceptible to oxidative damage (Weber et al., 2012). Studies on genes of the mammalian α7 family, which includes *Drosophila Dα6* gene, have been shown to play a role in neuroprotection by inducing the antioxidant system through Jak2/STAT3 pathway (Egea et al., 2015). Therefore, an absence of Dα6 subunits from neuronal membranes under conditions of spinosad exposure may increase susceptibility to oxidative damage.

Lysosomal dysfunction provides a parsimonious explanation as the cause for the mitochondrial impairment and ROS generated by spinosad exposure (Deus et al., 2020; Plotegher and Duchen, 2017; Stepien et al., 2020; Yambire et al., 2018) (**Figure 10**). But the accumulation of superoxide was observed earlier (1 hr) than the lysosomal defects (2 hr), although levels of Dα6 protein were shown to have decreased significantly after 30 min (**Figure 2 – figure supplement 2**). This could be explained by different capacities of DHE and LysoTracker to detect thresholds of damage that have a significant biological impact. However, it also leaves open the possibility that the generation of ROS is due to another mechanism that probably relates to the severe lowering of cardiolipins in mitochondrial membranes.

### Spinosad causes neurodegeneration and affects behavior in adults

Both LSD (Darios and Stevanin, 2020) and oxidative stress (Liu et al., 2017; Martelli et al., 2020) can cause neurodegeneration. The evidence for spinosad-induced neurodegeneration comes from the reduced climbing ability caused by chronic low dose exposures (McGurk et al., 2015; **Figure 7**), blindness (**Figure 8**), vacuolation of the lamina cartridges and severe vacuolation of adult CNS (**Figure 9**). Electroretinograms reveal that both *Dα6 knockout* mutants non-exposed and wild type flies chronically exposed to 0.2 ppm spinosad have reduced on-transients and amplitudes in response to light flashes (**Figure 8; Figure 9 – figure supplement 1**). *Dα6 knockout* mutants, however, show no vacuolation of lamina (**Figure 9 – figure supplement 1**). Given that *Dα6 knockout* mutants are viable, highly resistant to spinosad and show no conspicuous behavioral defects, it becomes clear that the majority of the impacts caused by spinosad are not initiated by the absence of Dα6 from neuronal membranes. The astonishing level of neurodegeneration observed in the central brain (**Figure 9G, H**) seems to be largely contained to the functional regions of the optic tubercle, mushroom body and superior lateral and medial protocerebrum. These regions are important centres for vision and memory, and learning and cognition in flies (Schürmann, 2016). Neurodegeneration in these regions indicate that a wide range of behaviours would be critically compromised in exposed flies.

Da6 nAChRs are not known to be expressed in photoreceptor cells or glial cells, but their expression in lamina (**Figure 8 – figure supplement 1**) supports their presence in post-synaptic cells. The accumulation of LD in PR after spinosad exposure (**Figure 8A**) suggests the existence of cell non-autonomous mechanisms initiated by spinosad in post-synaptic cells. Liu et al. (2017) showed that ROS induce the formation of lipids in neurons that are transported to glia, where they form LD. Here, a ROS signal generated by spinosad exposure in post-synaptic cells might be carried to PR, affecting lipid metabolism, and triggering LD accumulation. This hypothesis needs further investigation.

### Rational control of insecticide usage

In the public domain, organic insecticides are often assumed to be safer than synthetic ones for the environment and non-target insect species. The synthetic insecticide, imidacloprid, has faced intense scrutiny and bans because of its impact on the behavior of bees and the potential for this to contribute to the colony collapse phenomenon (Wu-Smart and Spivak, 2016). No other insecticide has been so comprehensively investigated, so it is not yet clear whether other chemicals pose similar risks. This study has revealed disturbing impacts of low doses of an organic insecticide, spinosad. Using the same methods deployed here, imidacloprid had a lower impact in *Drosophila* than spinosad (Martelli et al., 2020). At the same low acute dose (2.5 ppm for 2 hr), imidacloprid has no impact on larval survival, while spinosad is lethal. 4 ppm imidacloprid causes blindness and neurodegeneration, but no brain vacuolation under conditions of chronic exposure with 56% of flies dying in 25 days. 0.2ppm spinosad causes blindness and widespread brain vacuolation with 54% of flies dying in 25 days. That the nAChR Dα6 subunit has been shown to be a highly conserved spinosad target across a wide range of insects (Perry et al., 2015) suggests that low doses of this insecticide may have similar impacts in other species. The susceptibility of different species to insecticides varies, so the doses required may differ between them. The protocols used here will be useful in assessing the risk that spinosad poses to beneficial insects. Given the extent to which spinosad affects mitochondrial function, lipid metabolism and the brain, this insecticide may compromise the capacity of insects to survive in natural populations exposed to a variety of stresses including some of those that are being linked to insect population declines (Cardoso et al., 2020; Sánchez-Bayo and Wyckhuys, 2019).

Two clocks are ticking. The global human population is increasing and the amount of arable land available for food production is decreasing. Thus, the amount of food produced per hectare needs to increase. Our capacity to produce enough food has been underpinned by the use of insecticides. Approximately 600,000 tonnes of insecticides are used annually around the world (Aizen et al., 2009; Klein et al., 2007), but sublethal concentrations found in contaminated environments can affect behaviour, fitness and development of target and non-target insects (Müller, 2018). Despite their distinct modes of action, spinosad and imidacloprid produce a similar spectrum of damage (Martelli et al., 2020). This similarity arises because both insecticides trigger oxidative stress in the brain, albeit via different mechanisms. Several other insecticide classes such as organochlorines, organophosphates, carbamates and pyrethroids have all been shown to promote oxidative stress (Balieira et al., 2018; Karami-Mohajeri and Abdollahi, 2011; Lukaszewicz-Hussain, 2010; Terhzaz et al., 2015; Wang et al., 2016). Many insect populations are exposed to a continuously changing cocktail of insecticides (Kerr, 2017; Tosi et al., 2018), most of which are capable of producing ROS. The cumulative impact of these different insecticides could be significant. Our research clarifies the mode of action of spinosad, highlighting the perturbations and damage that occur downstream of the insecticide:receptor interaction. Other chemicals should not be assumed to be environmentally safe until their low dose biological impacts have been examined in similar detail.

## Material and Methods

### Fly strains and rearing

Armenia^14^ (Line 14), an isofemale line derived from Armenia^60^ (Drosophila Genomics Resource Center #103394) (Perry et al., 2008), was used as the susceptible wild type line for all assays except the following. Expression of nAChR-*Dα6* gene in adult brains: *Dα6* T2A Gal4 (BDSC #76137) was crossed with UAS-GFP.nls (BDSC #4775). Insecticide impact on mitochondrial turnover: the MitoTimer line (Gottlieb and Stotland, 2015) was used. GCaMP experiment: UAS-tdTomato-P2A-GCaMP5G (III) (Daniels et al., 2014; Wong et al., 2014) was crossed with *Dα6* T2A Gal4 (BDSC #76137). Two mutants for the nAChR-*Dα6* gene, which confers resistance to spinosad (Perry et al 2015) and their background control lines were used to investigate the insecticide mode of action. The first of these is Line 14 *Dα6 KO* strain, a mutant recovered following EMS mutagenesis in the Line 14 genetic background, with no detectable *Dα6* expression (Perry et al., 2015). The second mutant is a CRISPR knockout of *Dα6* generated in the CantonS genetic background. For experiments aiming to investigate the trafficking of *Dα6* nAChR in brains, UAS *Dα6* CFP tagged strain built in Line 14 *Dα6 KO* background (obtained by CRISPR) was crossed to a Gal4-L driver in Line 14 *Dα6 KO* background strain. For experiments involving larvae, flies were reared on standard food media sprinkled with dried yeast and maintained at 25°C. For experiments involving adults, flies were reared in molasses food and maintained at 25°C. In all experiments involving adult flies only females were used to maintain consistency.

### Insecticide dilution and exposure

The pure version of spinosad (Sigma Aldrich®) was used in all assays. The chemical was diluted to create 1000 ppm stocks solution, using dimethyl sulfoxide (DMSO), and was kept on freezer (−20°C). Before exposures, 5x stocks were generated for the dose being used by diluting the 1000 ppm stock in 5% Analytical Reagent Sucrose (Chem Supply) solution (or equivalent dose of DMSO for controls).

### Antioxidant treatment

The antioxidant, N-acetylcysteine amide (NACA) was used as previously described (Martelli et al., 2020). Briefly, larvae were treated with 300 µg/mL of NACA in 5% Analytical Reagent Sucrose (Chem Supply) solution for 5 hr prior to exposure to spinosad exposures.

### Fly medias used

| Standard Food (1L) |  | Apple Juice Plates (1L) |  | Molasses Food (1L) |  |
| --- | --- | --- | --- | --- | --- |
| H<sub>2</sub>O | 987 mL | H<sub>2</sub>O | 720 mL | H<sub>2</sub>O | 800 mL |
| Potassium Tartrate | 8.0 g | Agar | 20 g | Molasses | 160 mL |
| Calcium Chloride | 0.5 g | Apple Juice | 200 mL | Maize meal | 60 g |
| Agar | 5.0 g | Brewer's Yeast | 7.0 g | Dried active yeast | 15 g |
| yeast | 12 g | Glucose | 52 g | Agar | 6.0 g |
| Glucose | 53 g | Sucrose | 26 g | Acid mix | 7.5 mL |
| Sucrose | 27 g | Tegosept | 6.0 mL | Tegosept | 5.0 mL |
| Semolina | 67 g |  |  |  |  |
| Acid Mix | 12 mL |  |  |  |  |
| Tegosept | 15 mL |  |  |  |  |

### Larvae movement assay

Larvae movement in response to insecticide exposure was quantified by Wiggle Index Assay, as described by Denecke et al. (2015). 25 third instar larvae were used for a single biological replicate and four replicates were tested for each exposure condition. Undosed larvae in NUNC cell plates (Thermo-Scientific) in 5% Analytical Reagent Sucrose (Chem Supply) solution were filmed for 30 seconds and then 30 min, 1 hr, 1 hr and 30 min and 2 hr after spinosad exposure. The motility at each time-point is expressed in terms of Relative Movement Ratio (RMR), normalized to motility prior to spinosad addition.

### Larvae viability and adult survival tests

For all tests 5 replicates of 20 individuals (100 individuals) per condition were used. In assessing third instar larval viability and metamorphosis following insecticide exposure, individuals were rinsed three times with 5% w/v sucrose (Chem Supply) and placed in vials on insecticide-free food medium. Survival probability of larvae exposed to 2.5 ppm spinosad for 2 hr was analysed using Kaplan-Meier method and the Log-rank Mantel-Cox test. Correct percentage survival of larvae exposed to 0.5 ppm spinosad for 2 hr, or 0.1 ppm spinosad for 4 hr was analysed using Abbots’ correction. To examine the survival of adult flies chronically exposed to 0.2 ppm spinosad, 5 replicates of 20 females (3-5 days old) were exposed for 25 days. The same number of flies was used for the control group. Statistical analysis was based on the Kaplan-Meier method and data were compared by the Log-rank Mantel-Cox test.

### GCaMP assay

Cytosolic [Ca2+] in *Drosophila* primary neurons was measured as previously described (Martelli et al., 2020). Briefly, four brains from third instar larvae were dissected to generate ideal number of cells for 3 plates. Cells were allowed to develop in culture plates (35 mm glass-bottom dishes with 10 mm bottom well (Cellvis), coated with concanavalin A (Sigma)) with Schneider’s media for 4 days with the media refreshed daily. Recording was done using a Nikon A1 confocal microscope, 40x air objective, sequential 488nm and 561nm excitation. Measurements were taken at 3 second intervals. Cytosolic Ca^2+^ levels were reported as GCaMP5G signal intensity divided by tdTomato signal intensity. Signal was recorded for 60 sec before the addition of 2.5 ppm or 25 ppm spinosad to the bath solution. 5 min after that, both insecticide and control groups were stimulated by the cholinergic agonist carbachol (100 µM) added to the bath solution, and finally the SERCA inhibitor thapsigargin (5 µM) was added after a further 1 min. At least 50 neuronal cells were evaluated per treatment. The data were analysed using a Student’s t-test.

### Evaluation of mitochondrial turnover

Mitochondrial turnover was assessed as previously described (Martelli et al., 2020). Larvae of the MitoTimer line were exposed to 2.5 ppm spinosad for 2 hr. Control larvae were exposed to 2.5ppm DMSO. Midguts and brains were dissected in PBS and fixed in 4% PFA (Electron Microscopy Science) and mounted in Vectashield (Vector Laboratories). 20 anterior midguts and 20 pairs of optical lobes were analysed for each condition. Confocal microscopy images were obtained in Leica SP5 Laser Scanning Confocal Microscope at 200x magnification for both green (excitation/emission 488/518 nm) and red (excitation/emission 543/572 nm) signals. Three independent measurements along the z stack were analysed for each sample. Fluorescence intensity was quantified on ImageJ software and data were analysed using a Student’s t-test.

### Systemic mitochondrial aconitase activity

Relative mitochondrial aconitase activity was quantified using the colorimetric Aconitase Activity Assay Kit from Sigma (#MAK051), following manufacturer’s instructions as previously described (Martelli et al., 2020). A total of six biological replicates (25 whole larvae per replicate) were exposed to 2.5 ppm spinosad for 2 hr, whilst six control replicates (25 whole larvae per replicate) were exposed to DMSO for 2 hr. Absorbance was measured at 450 nm in a FLUOstar OPTIMA (BMG Labtech) microplate reader using the software OPTIMA and normalized to sample weight. The data were analysed using a Student’s t-test.

### Systemic ATP levels

Relative ATP levels were quantified fluorometrically using an ATP assay kit (Abcam, #83355), following manufacturer instructions as previously described (Martelli et al., 2020). A total of six biological replicates (20 larvae per replicate) were exposed to 2.5 ppm spinosad for 2 hr, whilst six control replicates (20 larvae per replicate) were exposed to DMSO for 2 hr. Fluorescence was measured at excitation/emission = 535/587 nm in FLUOstar OPTIMA (BMG Labtech) microplate reader using the software OPTIMA and normalized to sample weight. The data were analysed using a Student’s t-test.

### Measurement of superoxide (O_2_^-^) levels

Levels of superoxide were assessed dihydroethidium staining (DHE – Sigma-Aldrich), as described in (Owusu-Ansah et al. 2008). Briefly, larvae were dissected in Schneider’s media (GIBCO) and incubated with DHE at room temperature on an orbital shaker for 7 minutes in dark. Tissues were fixed in 8% PFA (Electron Microscopy Science) for 5 minutes at room temperature on an orbital shaker in dark. Tissues were then rinsed with PBS (Ambion) and mounted in Vectashield (Vector Laboratories). Confocal microscopy images were obtained in a Leica SP5 Laser Scanning Confocal Microscope at 200x magnification (excitation/emission 518/605 nm). Third instar larvae were exposed to 2.5 ppm spinosad for 1 or 2 hr. Controls were exposed to equivalent doses of DMSO. A total of 15 brains and 15 midguts were assessed for each condition. Three independent measurements along the z stack were analysed for each sample. Fluorescence intensity was quantified on ImageJ software and data were analysed using a Student’s t-test.

### Evaluation of lipid environment of metabolic tissues in larvae

Fat bodies, midguts and Malpighian tubules were dissected in PBS (Ambion) and subjected to lipid staining with Nile Red N3013 Technical grade (Sigma-Aldrich) as previously described (Martelli et al., 2020). Three biological replicates were performed for each exposure condition, each replicate consisting of a single tissue from a single larva. Tissues were fixed in 4% PFA (Electron Microscopy Science) and stained with 0.5 µg/mL Nile Red/PBS for 20 minutes in dark. Slides were mounted in Vectashield (Vector Laboratories) and analysed using a Leica SP5 Laser Scanning Confocal Microscope at 400x magnification. Red emission was observed with 540 ± 12.5 nm excitation and 590 LP nm emission filters. Images were analysed using ImageJ software. For fat bodies, the number, size and percentage of area occupied by lipid droplets was measure in 5 different random sections of 2500 µm^2^ per sample (three samples per group). For Malpighian tubules number of lipid droplets was measure in five different random sections of 900 µm^2^ per sample (three samples per group). For midgut samples, lipid droplets were not quantified, rather zones containing lipid droplets were identified by microscopy. The data were analysed using Student’s t-test.

### Lipid quantification in larvae hemolymph

Extracted hemolymph lipids were measured using the sulfo-phospho-vanillin method (Cheng et al. 2011) as previously described (Martelli et al., 2020). 30 third instar larvae were used for a single biological replicate and 7 replicate samples were prepared for each exposure condition. Absorbance was measured at 540 nm in a CLARIOstar® (BMG LABTECH) microplate reader using MARS Data Analysis Software (version 3.10 R3). Cholesterol (Sigma-Aldrich) was used for the preparation of standard curves. The data were analysed using a Student’s t-test.

### Lipid Extraction and Analysis Using Liquid Chromatography-Mass Spectrometry

Lipidomic analyses of whole larvae exposed for 2 h to 2.5 ppm spinosad were performed in biological triplicate and analyzed by electrospray ionization-mass spectrometry (ESI-MS) using an Agilent Triple Quad 6410 as previously described (Martelli et al., 2020). Briefly, samples were transferred to CryoMill tubes treated with 0.001% BHT (butylated hydroxytoluene) and frozen in liquid nitrogen. Samples were subsequently homogenized using a CryoMill (Bertin Technologies) at −10 °C. Then 400 μL of chloroform was added to each tube and samples were incubated for 15 min at room temperature in a shaker at 1,200 rpm. Samples were then centrifuged for 15 min, at 13,000 rpm at room temperature; the supernatants were removed and transferred to new 1.5-mL microtubes. For a second wash, 100 μL of methanol (0.001% BHT and 0.01 g/mL 13C5 valine) and 200 μL of chloroform were added to CryoMill tubes, followed by vortexing and centrifugation as before. Supernatants were transferred to the previous 1.5-mL microtubes. A total of 300 μL of 0.1 M HCl was added to pooled supernatants and microtubes were then vortexed and centrifuged (15 min, room temperature, 13,000 rpm). Upper phases (lipid phases) were collected and transferred to clean 1.5-mL microtubes, as well as the lower phases (polar phases). All samples were kept at −20 °C until analysis. For liquid chromatography-mass spectrometry (LC-MS) analysis, microtubes were shaken for 30 min at 30 °C, then centrifuged at 100 rpm for 10 min at room temperature after which the supernatants were transferred to LC vials. Extracts were used for lipid analysis. For statistical analysis the concentration of lipid compounds was initially normalized to sample weight. Principal Components Analysis (PCA) was calculated to verify the contribution of each lipid compound in the variance of each treatment. PCA was calculated using the first two principal component axes. To discriminate the impacts of spinosad on the accumulation of specific lipid compounds we performed a One-way ANOVA test with post-hoc Tukey’s HSD (p<0.05).

### Investigating impacts on lysosomes

To investigate spinosad impacts on lysosomes the LysoTracker staining was used on larval brains dissected from 3^rd^ instars. Larvae were exposed to 2.5 ppm spinosad for 1 hr or 2 hr, in the last case brains were assessed immediately after the 2 hr exposure or 6 hr after that. Larvae were dissected in PBS and tissue immediately transferred to PBS solution containing LysoTracker Red DND-99 (1:10,000) (Invitrogen) for 7 minutes. Tissues were then rinsed 3 times in PBS and slides were mounted for immediate microscopy 400x magnification (DsRed filter). A total of 7 brain samples were assessed per group, with 3 random different sections of 900 µm^2^ accounted per brain. To investigate the hypothesis of Dα6 nAChRs being endocytosed and digested by lysosomes after exposure to 2.5 ppm spinosad for 2 hr, brains from larvae obtained by crossing UAS *Dα6* CFP tagged in Line 14 *Dα6 KO* strain to Gal4-L driver in Line 14 *Dα6 KO* strain were also subjected to LysoTracker staining. Images were analysed using the software ImageJ and data were analysed using Student’s t-test.

### Electrophysiology of the retina

Amplitudes and on transients were assessed as previously described (Martelli et al., 2020). Briefly, adult flies were anesthetized and glued to a glass slide. A reference electrode was inserted in the back of the fly head and the recording electrode was placed on the corneal surface of the eye, both electrodes were filled with 100 mM NaCl. Flies were maintained in the darkness for at least 5 min prior to a series of 1 s flashes of white light delivered using a halogen lamp. During screening 8 to 10 flies per treatment group were tested. For a given fly, amplitude and on transient measurements were averaged based on the response to the 3 light flashes. Responses were recorded and analysed using AxoScope 8.1. The data were analysed using Student’s t-test.

### Nile red staining of adult retinas

For whole mount staining of fly adult retinas, heads were dissected in cold PBS (Ambion) and fixed in 37% formaldehyde overnight. Subsequently, the retinas were dissected and rinsed several times with 1× PBS and incubated for 15 minutes at 1:1000 dilution of PBS with 1 mg/ml Nile Red (Sigma).

Tissues were then rinsed with PBS and immediately mounted with Vectashield (Vector Labs) for same-day imaging. For checking the effects of chronic exposures 8 retinas from 8 adult female flies were analysed per condition (imidacloprid 4 ppm and control) per day (after 1, 5, 10, 15 and 20 days of exposure). Images were obtained with a Leica TCS SP8 (DM600 CS), software LAS X, 600x magnification, and analysed using ImageJ. The data were analysed using Student’s t-test.

### Expression of Dα6 nAChRs in brain

The expression patter of nAChR-Dα6 gene in adult brains was assessed in the crossing between Dα6 T2A Gal4 (BDSC #76137) and UAS-GFP.nls (BDSC #4775). Adult brains were fixed in 4% PFA (Electron Microscopy Science) in PBS for 20 minutes at room temperature. PFA was removed and tissues were washed 3 times in PBS. Samples were mounted in Vectashield (Vector Laboratories). Images were obtained with a Leica TCS SP8 (DM600 CS), software LAS X, 400x magnification, using GFP channel. Images were analysed using the software ImageJ.

### Adult brain histology (Hematoxylin & Eosin staining)

Adult fly heads were fixed in 8% glutaraldehyde (EM grade) and embedded in paraffin. Sections (10 µm) were prepared by a microtome (Leica) and stained with Hematoxylin and Eosin as described (Chouhan et al., 2016). At least three animals were examined for each group (20 days exposure to 0.2 ppm spinosad plus control group) in terms of percentage of brain area vacuolated. The data were analysed using Student’s t-test.

### Transmission Electron Microscopy (TEM)

Laminas of adult flies chronically exposed to 0.2 ppm spinosad 20 days (controls exposed to equivalent volume of DMSO) were processed for TEM imaging as described (Luo et al., 2017). TEM of laminas of 20-day old CantonS and CantonS *Dα6 KO* mutants aged in the absence of spinosad was also investigated. Samples were processed using a Ted Pella Bio Wave microwave oven with vacuum attachment. Adult fly heads were dissected at 25 °C in 4 % paraformaldehyde, 2 % glutaraldehyde, and 0.1 M sodium cacodylate (pH 7.2). Samples were subsequently fixed at 4 °C for 48 hr. 1 % osmium tetroxide was used for secondary fixation with subsequent dehydration in ethanol and propylene oxide. Samples were then embedded in Embed-812 resin (Electron Microscopy Science, Hatfield, PA). 50 nm ultra-thin sections were obtained with a *Leica UC7* microtome and collected on Formvar-coated copper grids (Electron Microscopy Science, Hatfield, PA). Specimens were stained with 1 % uranyl acetate and 2.5 % lead citrate and imaged using a JEOL JEM 1010 transmission electron microscope with an AMT XR-16 mid-mount 16 mega-pixel CCD camera. For quantification of ultrastructural features, electron micrographs were examined from 3 different animals per treatment. The data were analysed using Student’s t-test.

### Bang Sensitivity

The bang sensitivity phenotype was tested after 1, 10 and 20 days of chronic exposure to 0.2 ppm spinosad. Flies were vortexed on a VWR vortex at maximum strength for 10 s. The time required for flies to flip over and regain normal standing posture was then recorded. The data were analysed using Wilcoxon signed-rank test.

### Climbing assay

Climbing phenotype was tested after 1, 10 and 20 days of exposure to 0.2 ppm spinosad. 5 adult female flies were placed into a clean vial and allowed to rest for 30 min. Vials were tapped against a pad and the time required for the flies to climb up to a pre-determined height (7 cm) was recorded.

Flies that did not climb the pre-determined height within 30 seconds were deemed to have failed the test. The data were analysed using Wilcoxon signed-rank test.

### Graphs and Statistical analysis

All graphs were created, and all statistical analysis were performed in the software R (v.3.4.3). Images were designed using the free image software Inkscape (0.92.4).

Many of the analyses performed here were conducted on spinosad and imidacloprid in parallel with these treatments sharing the same controls, allowing direct comparison of the impact of these insecticides. The imidacloprid data were published in (Martelli et al., 2020). The data with shared controls are shown in Fig 1 (A,D,E), Fig 3 (A,B,C,D,E,F), Fig 4 (A,B,C), Fig 4 - figure supplement 1, Fig 4 - figure supplement 2 (D,E), Fig 4 - figure supplement 3, Fig 4 - figure supplement 4, Fig 5 (E), Fig 6, Fig 6 - table supplement 1, Fig 7 (B) and Fig 8 (C,D,E).

## Funding

F.M. was supported by a Victorian Latin America Doctoral Scholarship, an Alfred Nicholas Fellowship, a UoM Faculty of Science Travelling Scholarship, and The Robert Johanson and Anne Swann Fund - Native Animals Trust (awarded to F.M. and T.P). P.B. was supported by the University of Melbourne. H.J.B. was supported by the Howard Hughes Medical Institute (HHMI) and is an investigator of HHMI. K.V. was supported by NIH (NIA) grant. Lipid analysis were performed at Metabolomics Australia at University of Melbourne, which is a National Collaborative Research Infrastructure Strategy initiative under Bioplatforms Australia Pty Ltd (http://www.bioplatforms.com/).

## Author Contribution

F.M., T.P., P.B. and H.J.B. conceived the study and designed the experiments. F.M. performed toxicology assays, all tissue confocal microscopy, behavioral assays, metabolic assays, and transcriptomics analysis. F.M. and Z.Z. processed and analysed the electron microscopy data. F.M. and J.W. performed the ERG analysis and RNAi experiments. F.M., T.R. and U.R. performed the lipidomic analysis. F.M., C.O.W., N.E.K. and K.V. performed GCaMP experiments. T.P., P.B. and H.J.B. acquired funding and supervised the research. F.M., P.B. and H.J.B. wrote the manuscript. All authors read, edited, reviewed, and approved the final version of the manuscript.

## Competing interests

The authors confirm that there are no competing interests.

## Supplementary Information for Martelli et al 2021

**Figure 2 - figure supplement 1.**
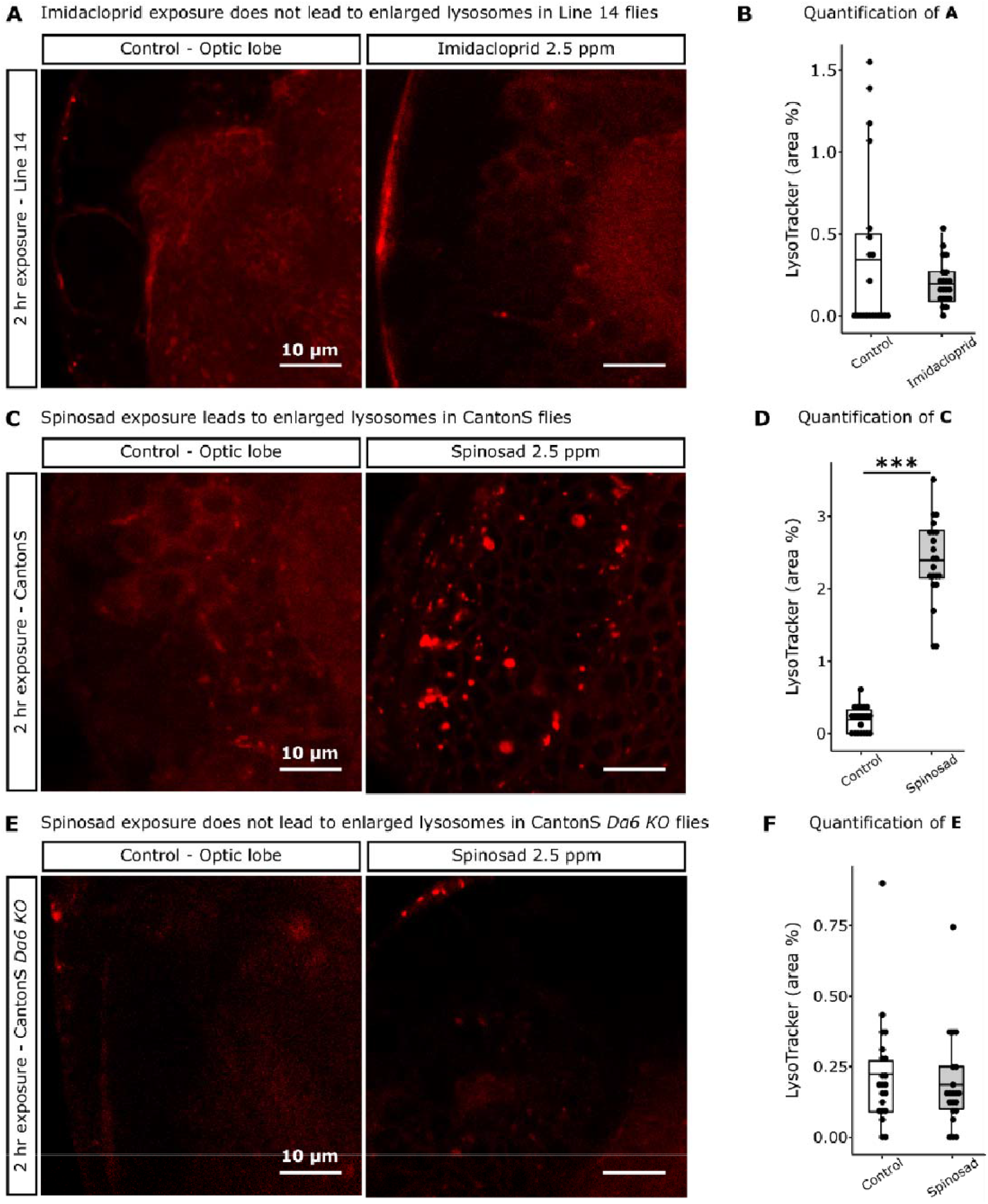
Enlarged lysosomes are only observed in response to spinosad exposure and in the presence of Dα6 nAChRs. **A**, Line 14 larvae exposed to 2.5 ppm imidacloprid for 2hr show no enlarged lysosomes in the brain. **B**, Quantification of A, LysoTracker area in the optic lobes (%) (n = 7 larvae/treatment, 3 optic lobe sections/larva). **C**, CantonS larvae exposed to 2.5 ppm spinosad for 2hr show significant increase in the number of enlarged lysosomes in the brain. **D**, Quantification of C, LysoTracker area in the optic lobes (%) (n = 7 larvae/treatment, 3 optic lobe sections/larva). **E**, CantonS *Dα6 knockout* larvae exposed to 2.5 ppm spinosad for 2hr show no enlarged lysosomes in the brain. **F**, Quantification of E, LysoTracker area in the optic lobes (%) (n = 7 larvae/treatment, 3 optic lobe sections/larva). Lysotracker staining, 400 x magnification. Microscopy images obtained in Leica SP5 Laser Scanning Confocal Microscope. t-test; ***P < 0.001.

**Figure 2 - figure supplement 2.**
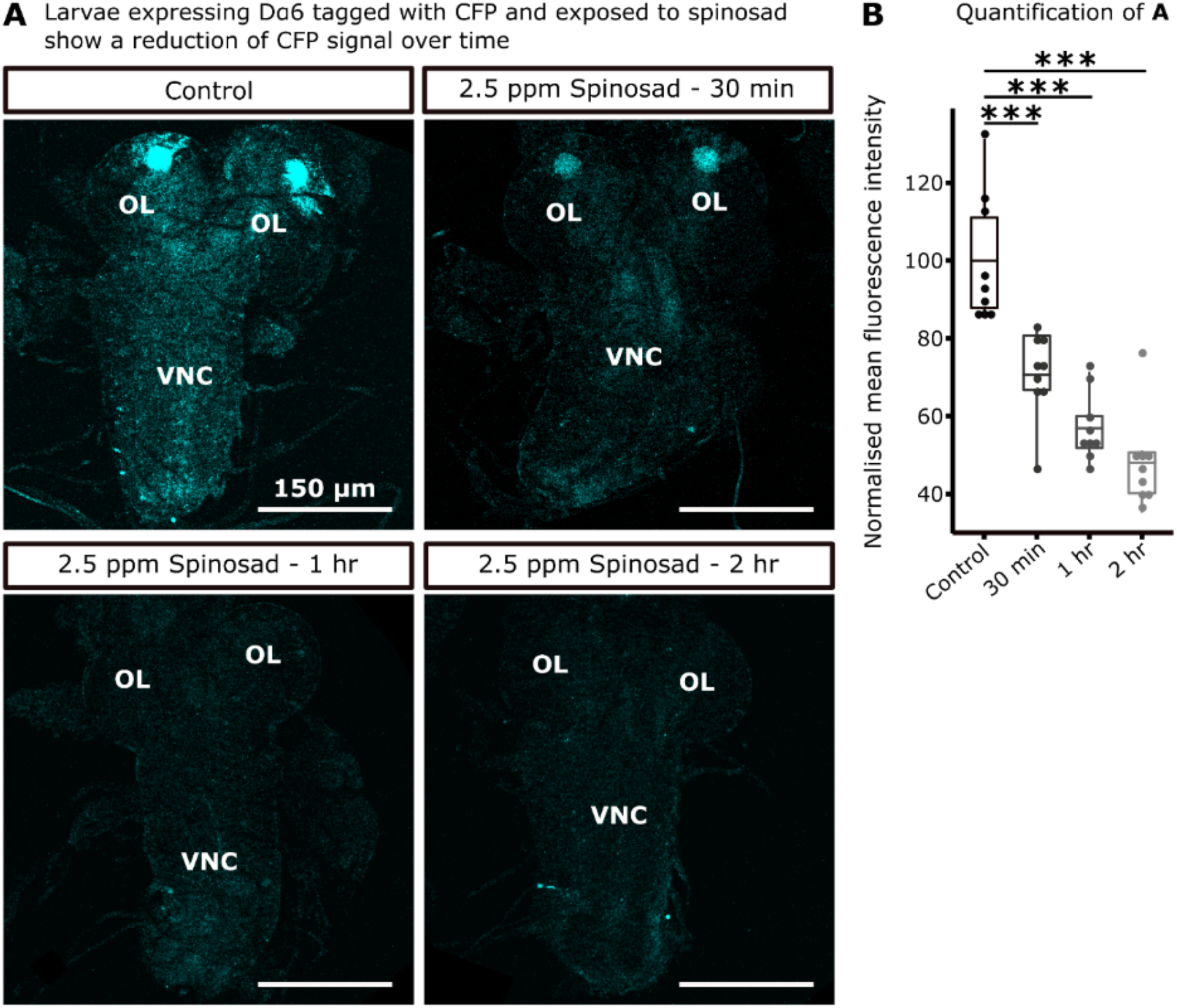
Exposure to spinosad reduces Dα6 nAChRs in neuronal membranes. **A**, Brains from larvae obtained by crossing UAS Dα6 CFP tagged in Line 14 *Dα6 KO* strain to Gal4-L driver in Line 14 *Dα6 KO* strain were exposed to 2.5 ppm spinosad for 30 min, 1 hr or 2 hr. **B**, Quantification of **A** (n = 3 larvae/condition, 3 brain sections/larva). Microscopy images obtained in Leica SP5 Laser Scanning Confocal Microscope, 400 x magnification. OP – optic lobe; VNC – ventral nerve cord. t-test; ***P < 0.001.

**Figure 4 - figure supplement 1.**
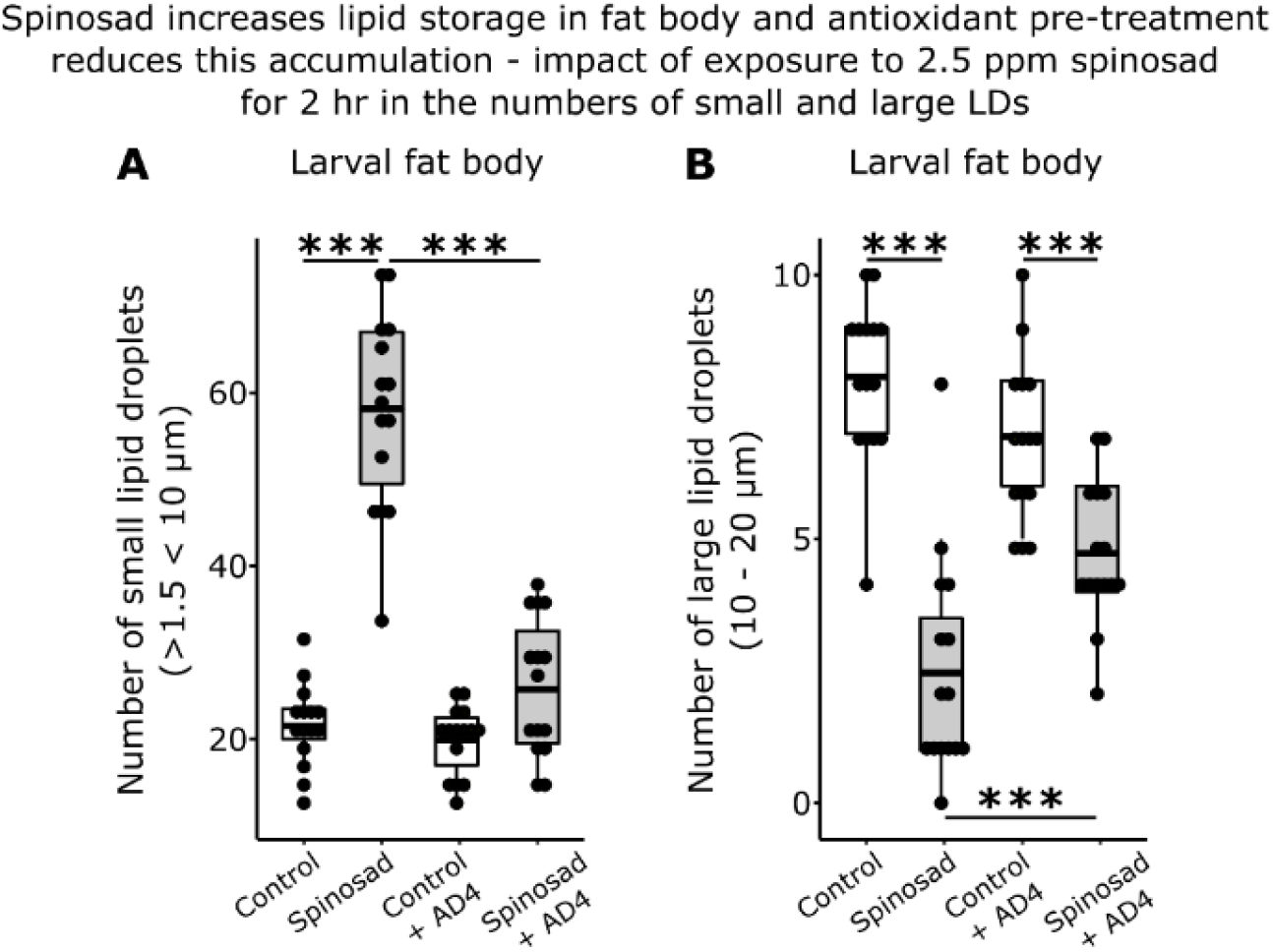
Impact of spinosad exposure on LD dynamics in fat body. Larvae exposed to 2.5 ppm spinosad for 2 hr show an accumulation of small LDs and reduction of large LD in the fat body. 5 hr pre-treatment with 300 µg/mL of antioxidant N-acetylcysteine amide (NACA) reduces this effect. **A**, Number of small LD (> 1.5 µm < 10 µm). **B**, Number of large LD (10 µm - 20 µm). n = 3 larvae/group; 5 image sections/larva. t-test; ***P < 0.001.

**Figure 4 - figure supplement 2.**
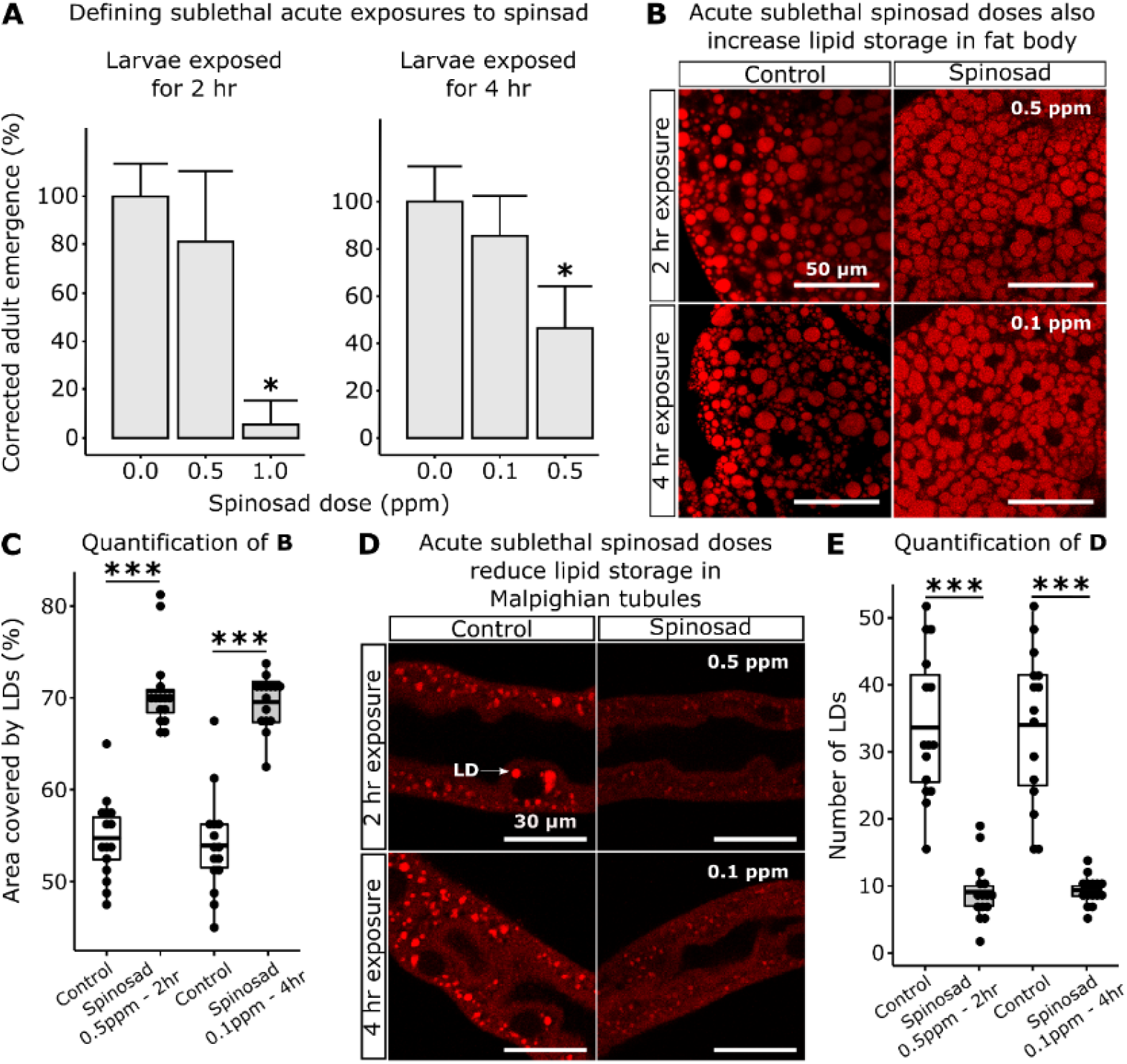
Spinosad doses that do not affect survival impact the larval lipid environment. **A**, Corrected adult emergence relative to controls - larvae exposed to different spinosad doses were rinsed in 5% sucrose and placed back onto insecticide-free media for quantification of adult emergence. 0.5 ppm for 2 hr and 0.1 ppm for 4 hr were determined as the highest doses that do not affect survival. **B**, Accumulation of LD in the fat body of larvae in response to the highest doses that do not affect survival. **C**, Percentage of area occupied by LD in fat body (n = 3 larvae/treatment; 5 image sections/larva). **D**, Reduction of lipid storage in Malpighian tubules of larvae exposed to the highest doses that do not affect survival. White arrow indicates a LD. **E**, Number of lipid droplets per Malpighian Tubule (n = 3 larvae/treatment; 5 sections/larva). Microscopy images obtained in Leica SP5 Laser Scanning Confocal Microscope, 400x magnification, Nile red staining. Error bars in **A** indicate 95% confidence interval (One-way ANOVA, Turkey’s HSD; *P < 0.05). **C** and **E**, t-test; ***P < 0.001.

**Figure 4 - figure supplement 3.**
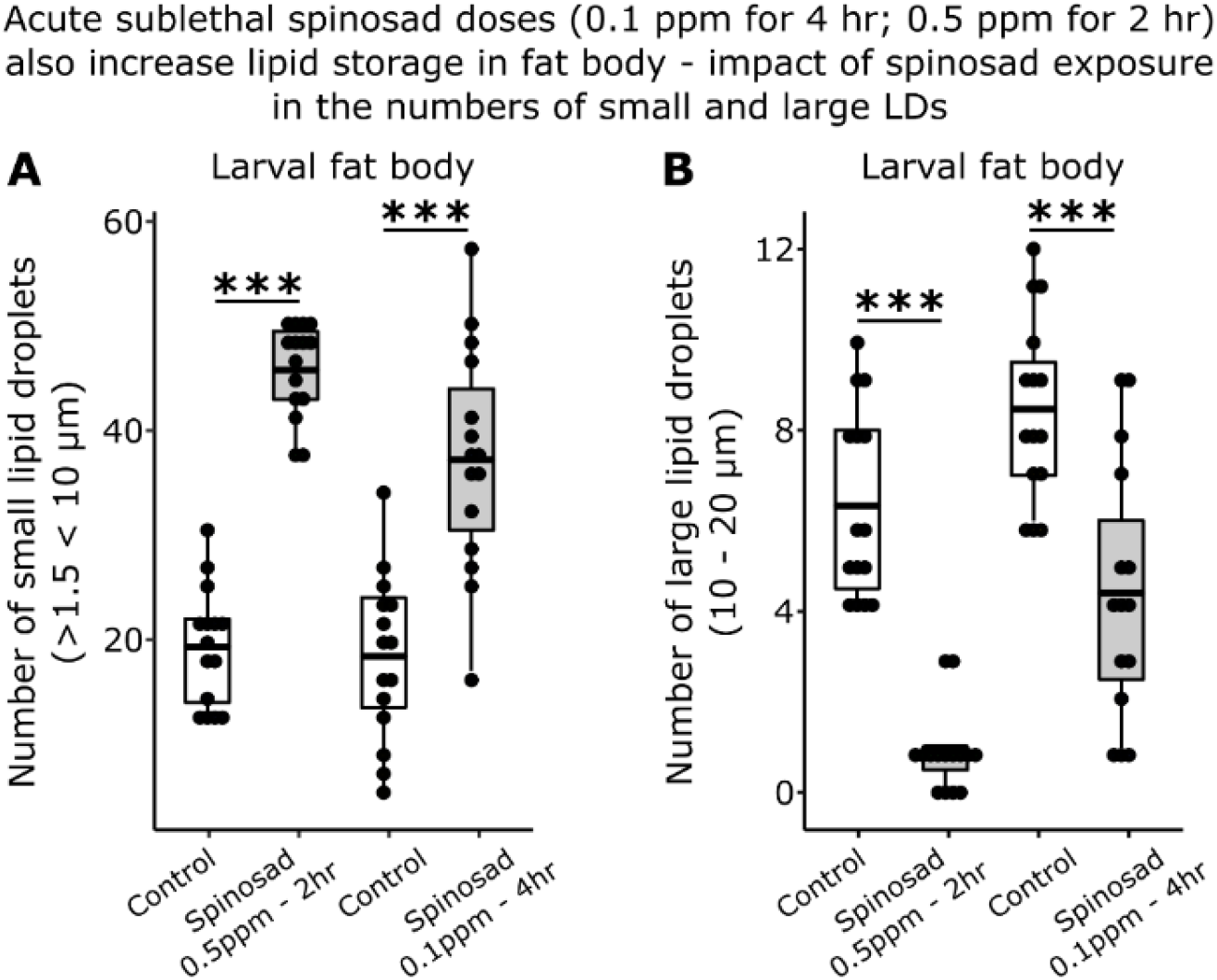
The highest spinosad doses that do not affect survival also impact LD dynamics in fat body. Larvae exposed to 0.5 ppm spinosad for 2 hr, or 0.1 ppm spinosad for 4 hr, show an accumulation of small LD and reduction of large LD in the fat body. **A**, Number of small LD (> 1.5 µm < 10 µm). **B**, Number of large LD (10 µm - 20 µm). n = 3 larvae/group; 5 image sections/larva. t-test; ***P < 0.001.

**Figure 4 - figure supplement 4.**
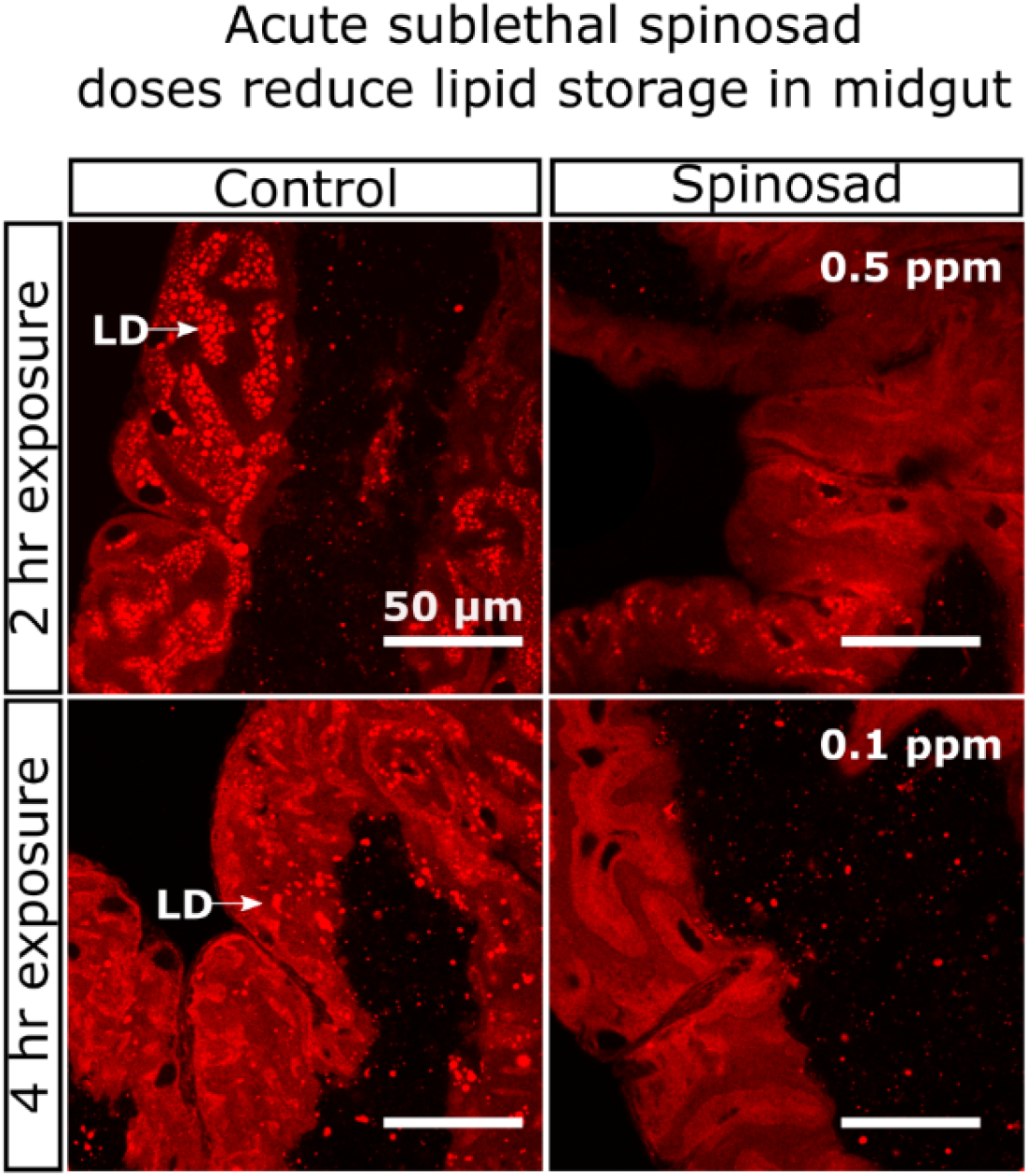
Spinosad doses that do not affect survival impact the larval lipid environment. Posterior midgut. White arrow indicates a cluster of LDs. Zones with LD accumulation were not quantified since they were only found in non-exposed animals (n = 3 larvae/ treatment). Microscopy images obtained in Leica SP5 Laser Scanning Confocal Microscope, 400x magnification, Nile red staining.

**Figure 8 - figure supplement 1.**
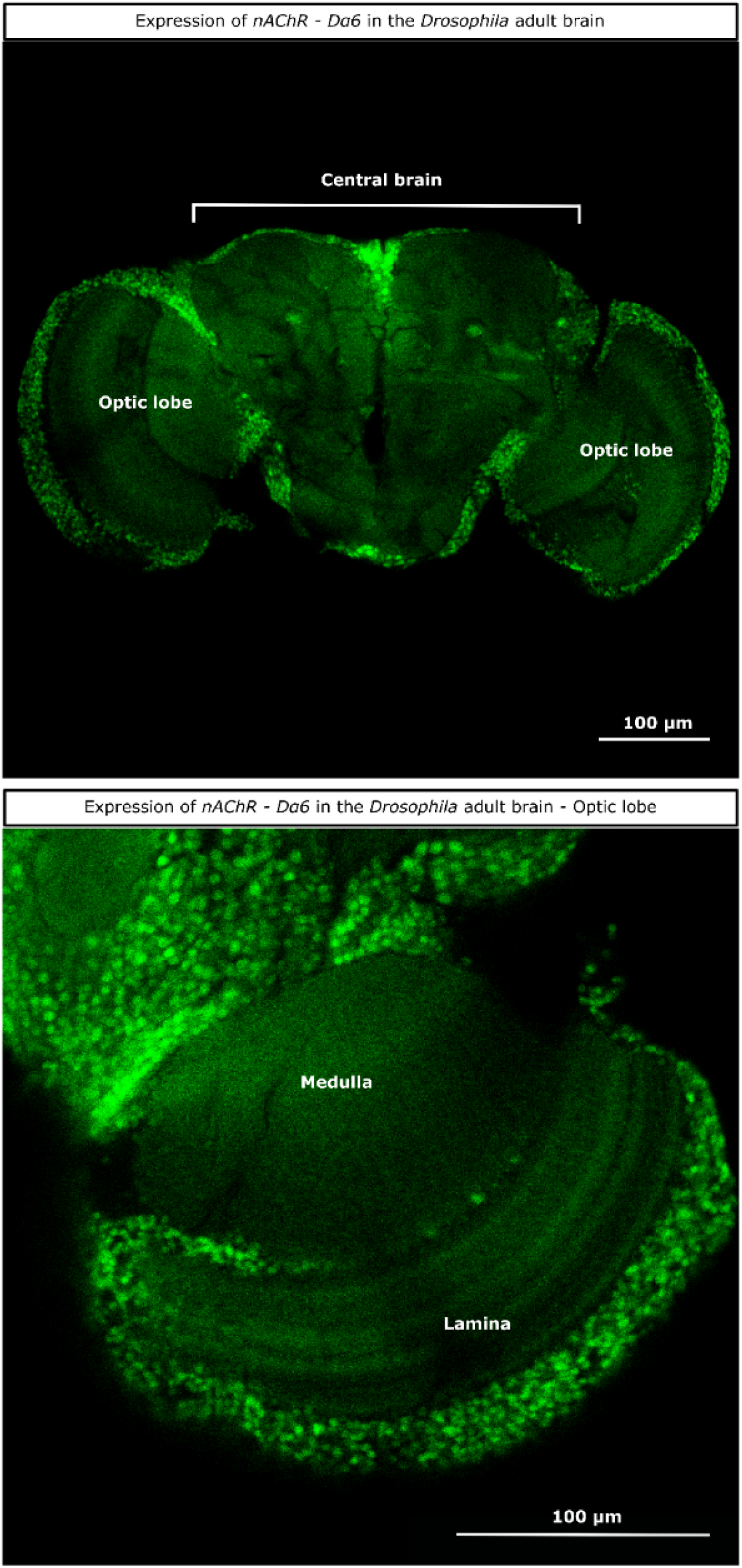
Expression pattern of nAChR subunit *Dα6* in the *Drosophila* adult brain. (*Dα6* T2A Gal4 > UAS-GFP.nls). Detail of the expression in lamina and medulla (optic lobe). Microscopy images obtained in Leica SP5 Laser Scanning Confocal Microscope. 400 x magnification.

**Figure 9 - figure supplement 1.**
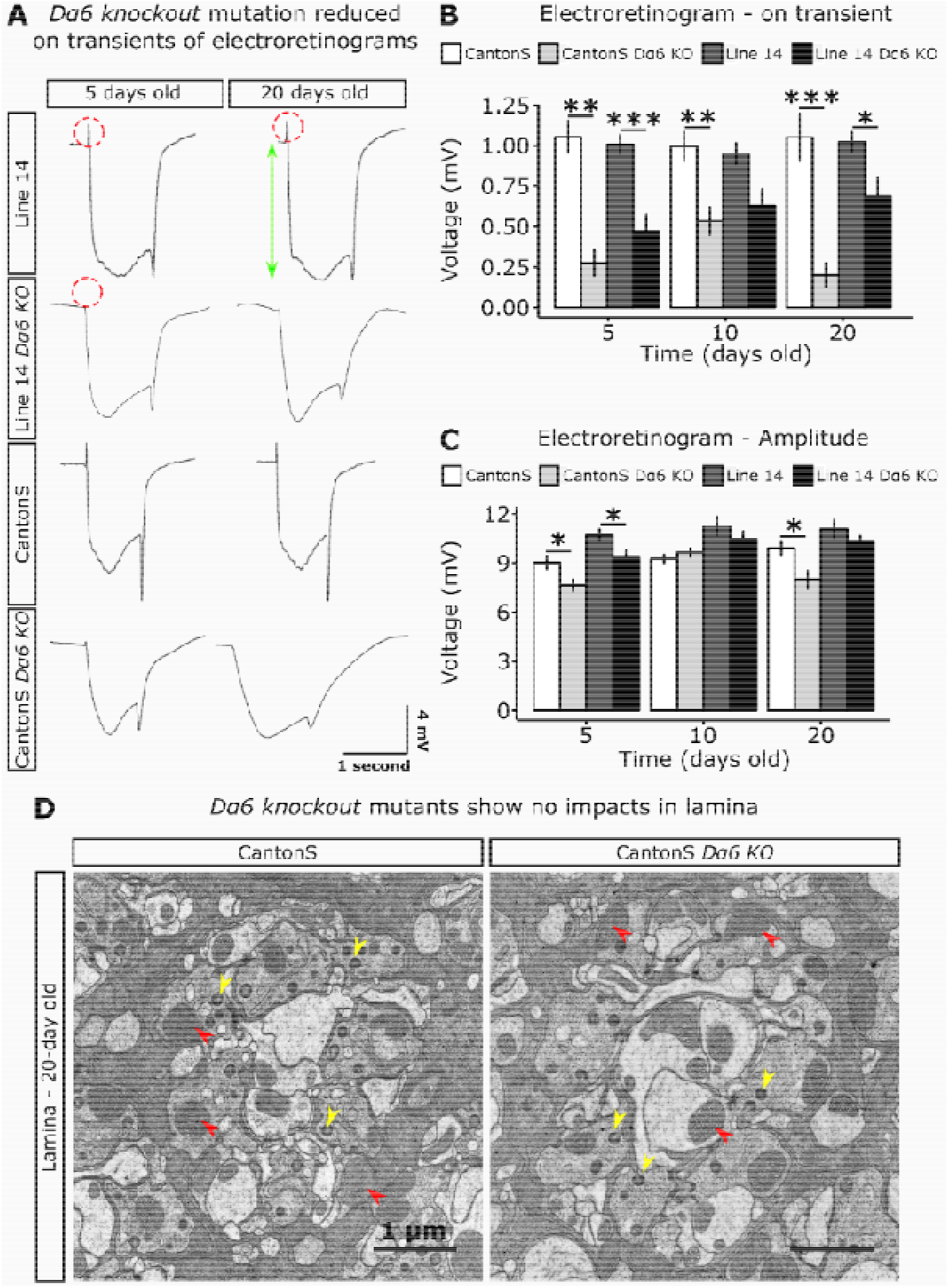
nAChR *Dα6 knockout* (*KO*) mutants show defective electroretinograms (ERGs) but no damage in lamina. **A**, ERGs of 5- and 20-days old females from Line 14, Line 14 *Dα6 KO* mutant, Canton S and Canton S *Dα6 KO* mutant. Red dotted circles indicate the on-transient signal and green arrow indicates the amplitude (n = 8 to 10 adult flies/strain/time point) **B**, On-transient signal of ERGs of 5-, 10- and 20-days old flies. **C**, Amplitude of ERGs of 5-, 10- and 20-days old flies. **D**, Electron microscopy of the lamina of 20-day old Canton S and Canton S *Dα6 KO* mutant flies aged in the absence of spinosad. Red arrowheads indicate normal mitochondria, yellow arrowheads indicate capitate projections. No conspicuous difference was noticed between mutant and background strains (10 images/fly; 3 flies/genotype). t-test; *P < 0.05, **P < 0.01, ***P < 0.001.

**Figure 6 - table supplement 1.**
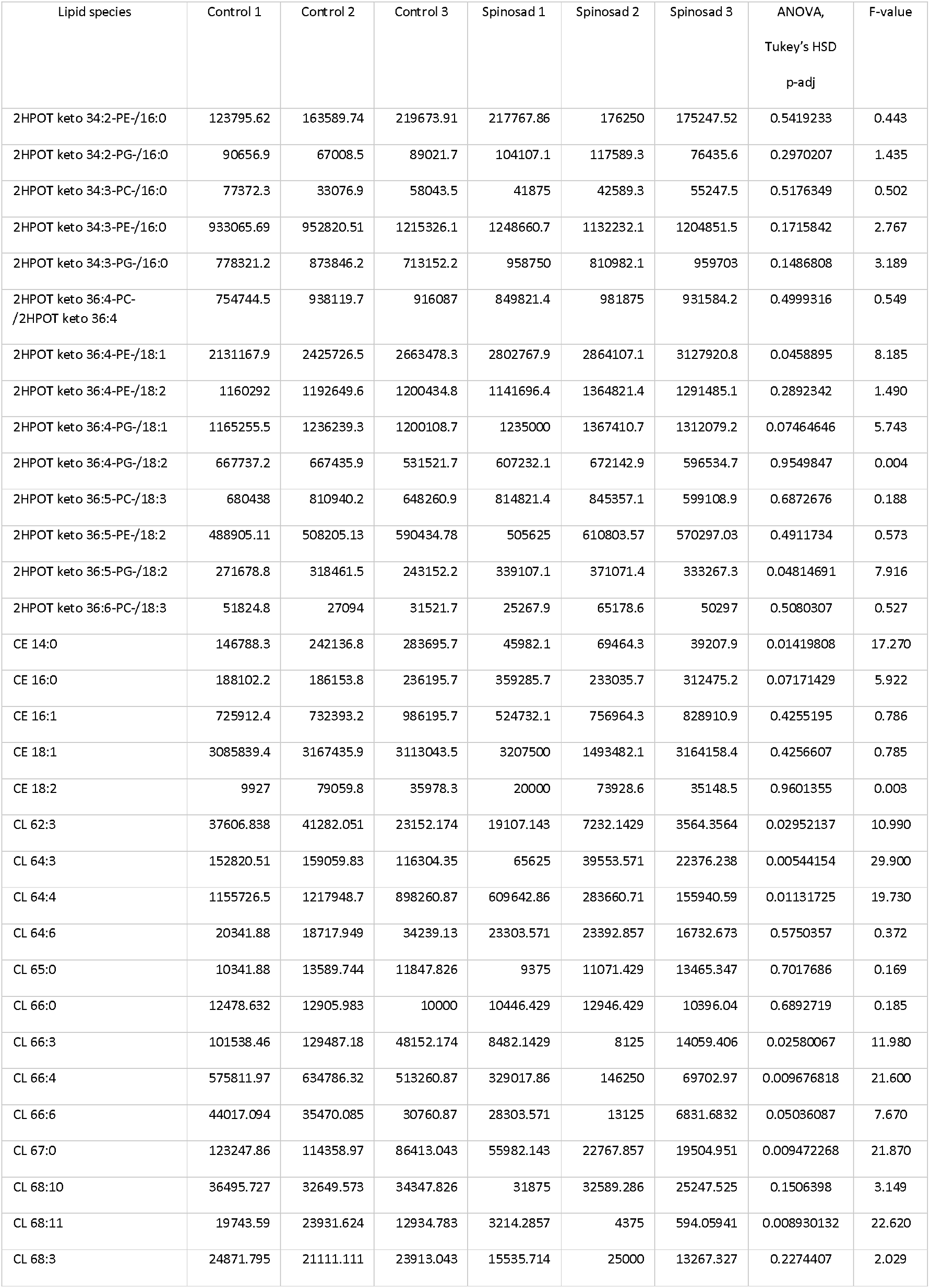
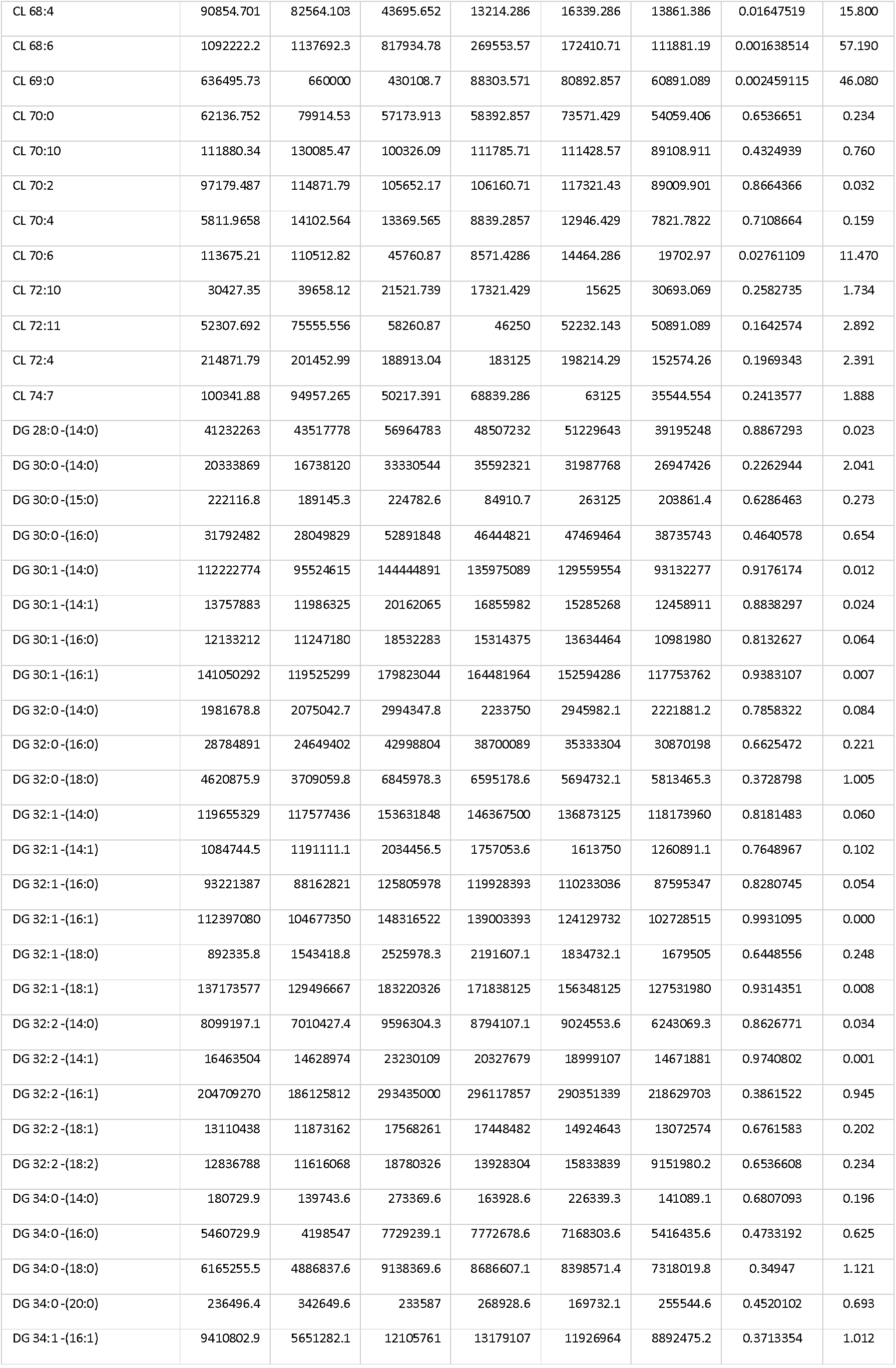
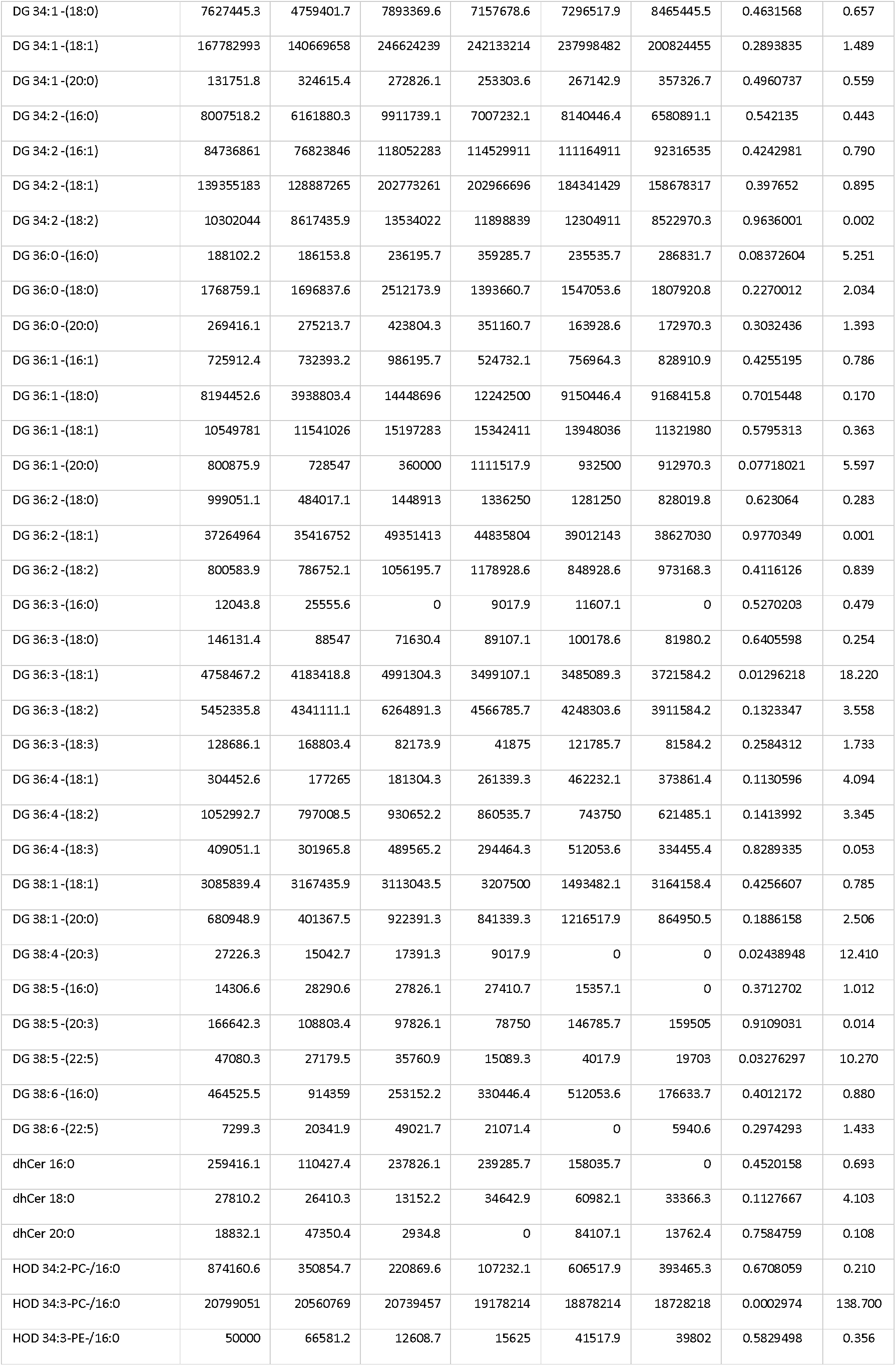
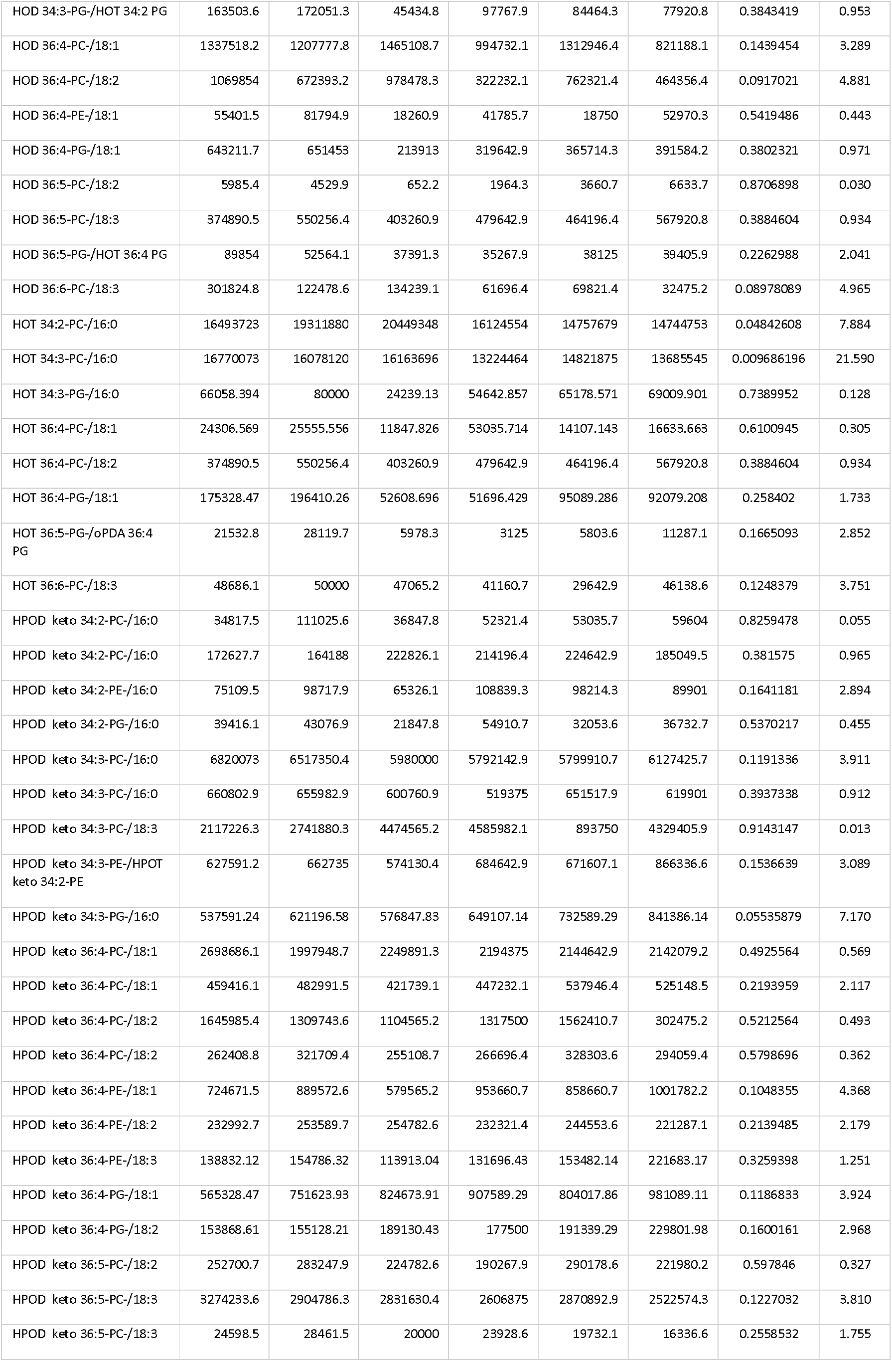
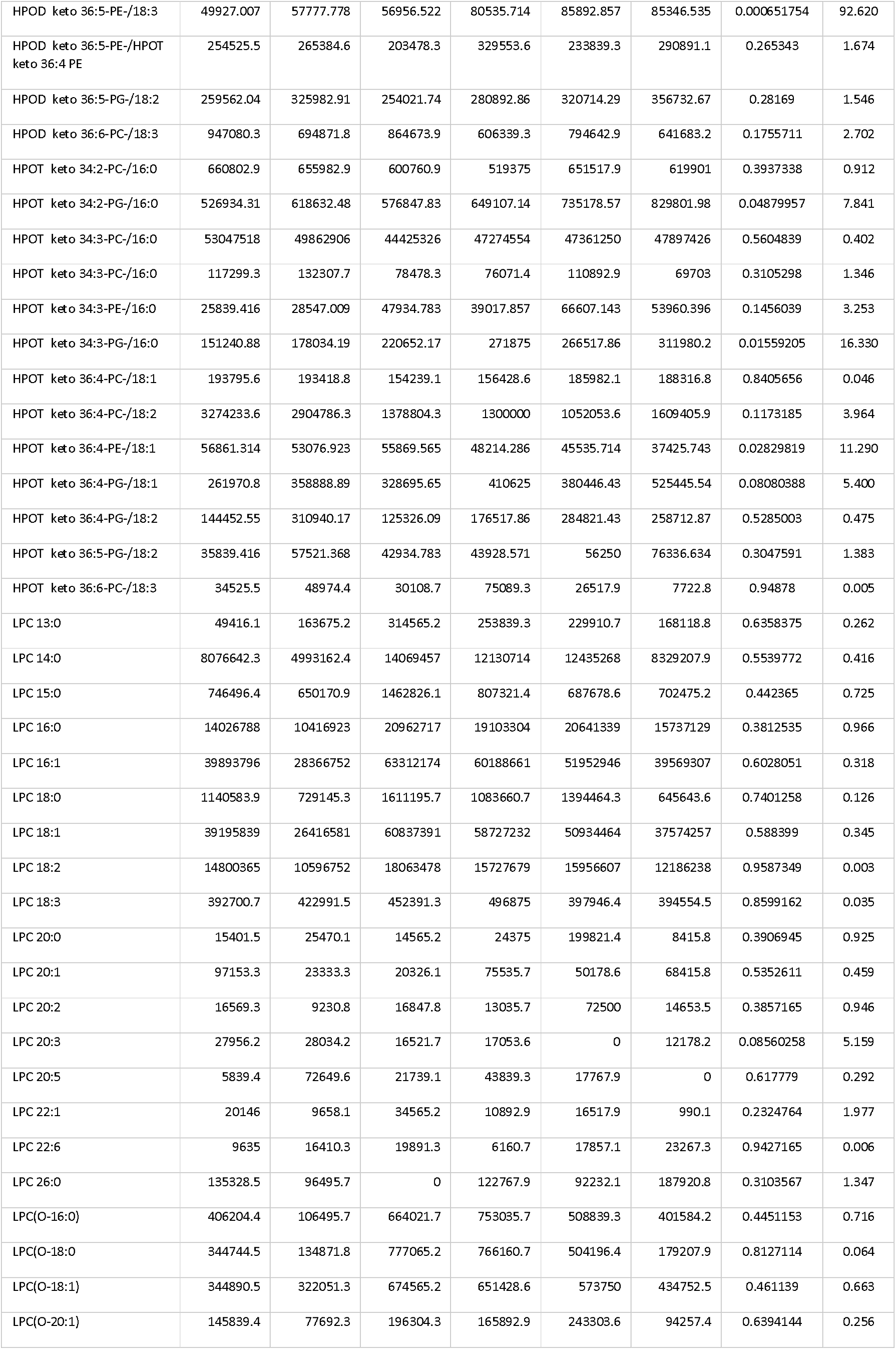
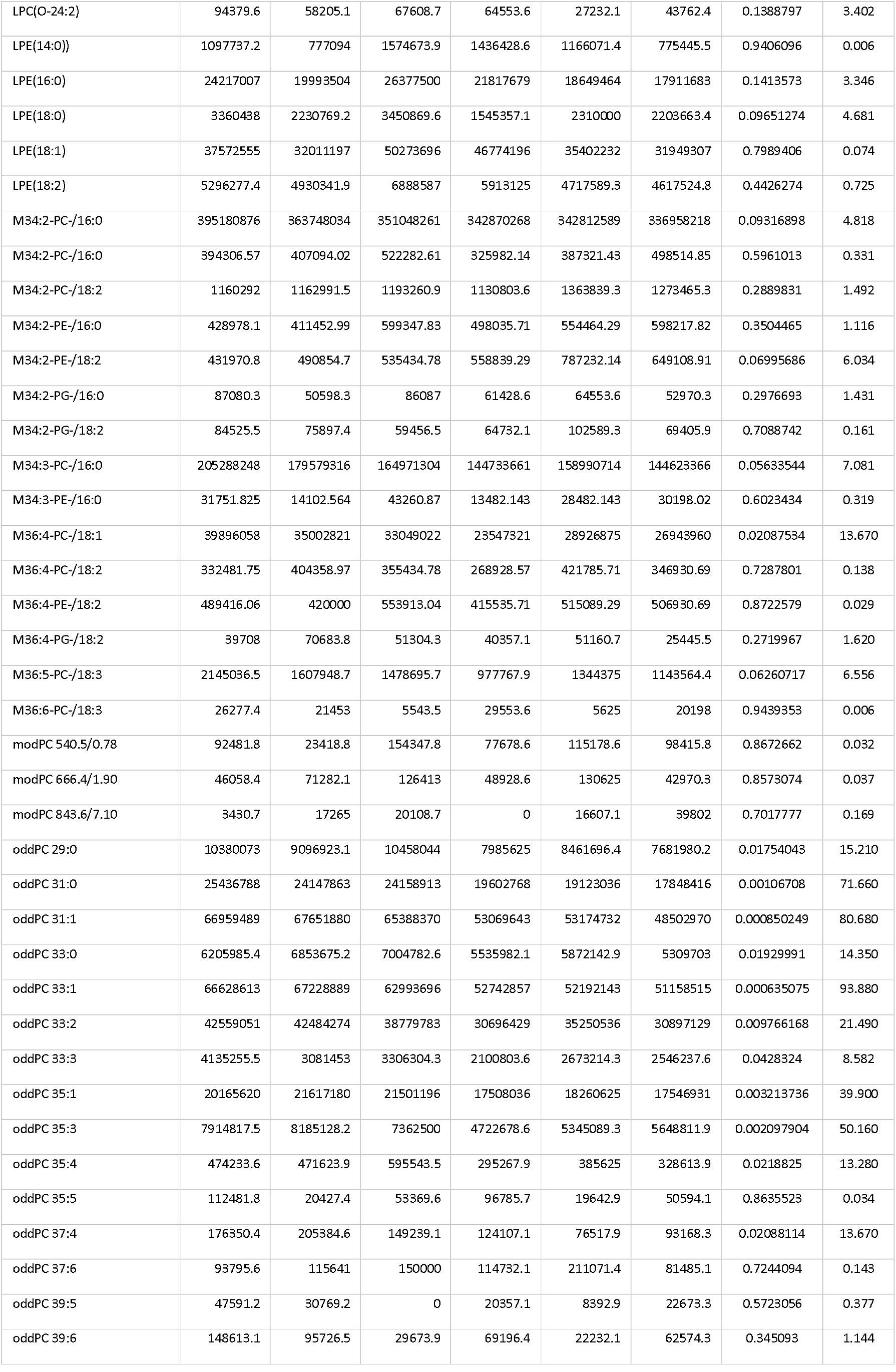
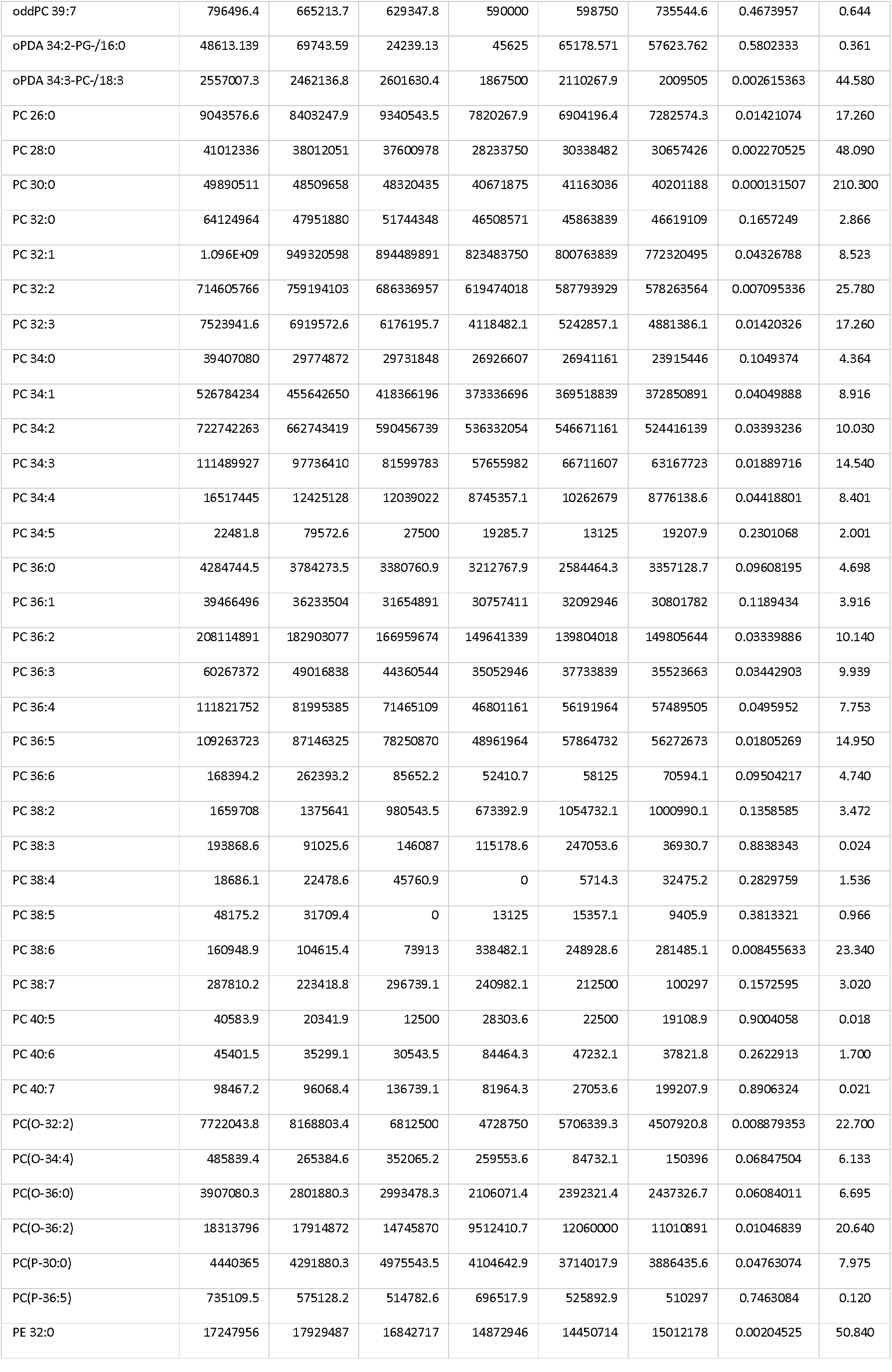
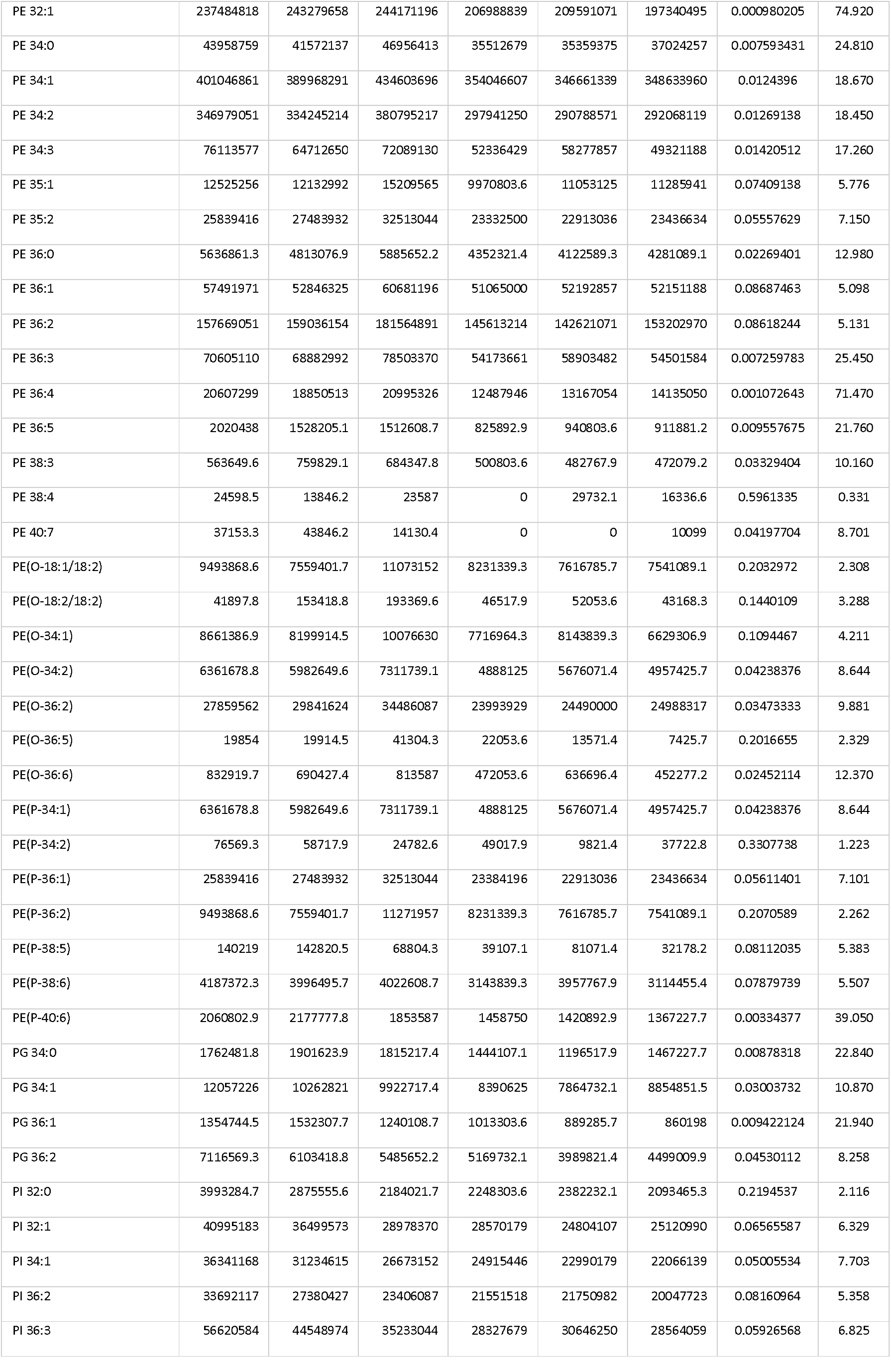
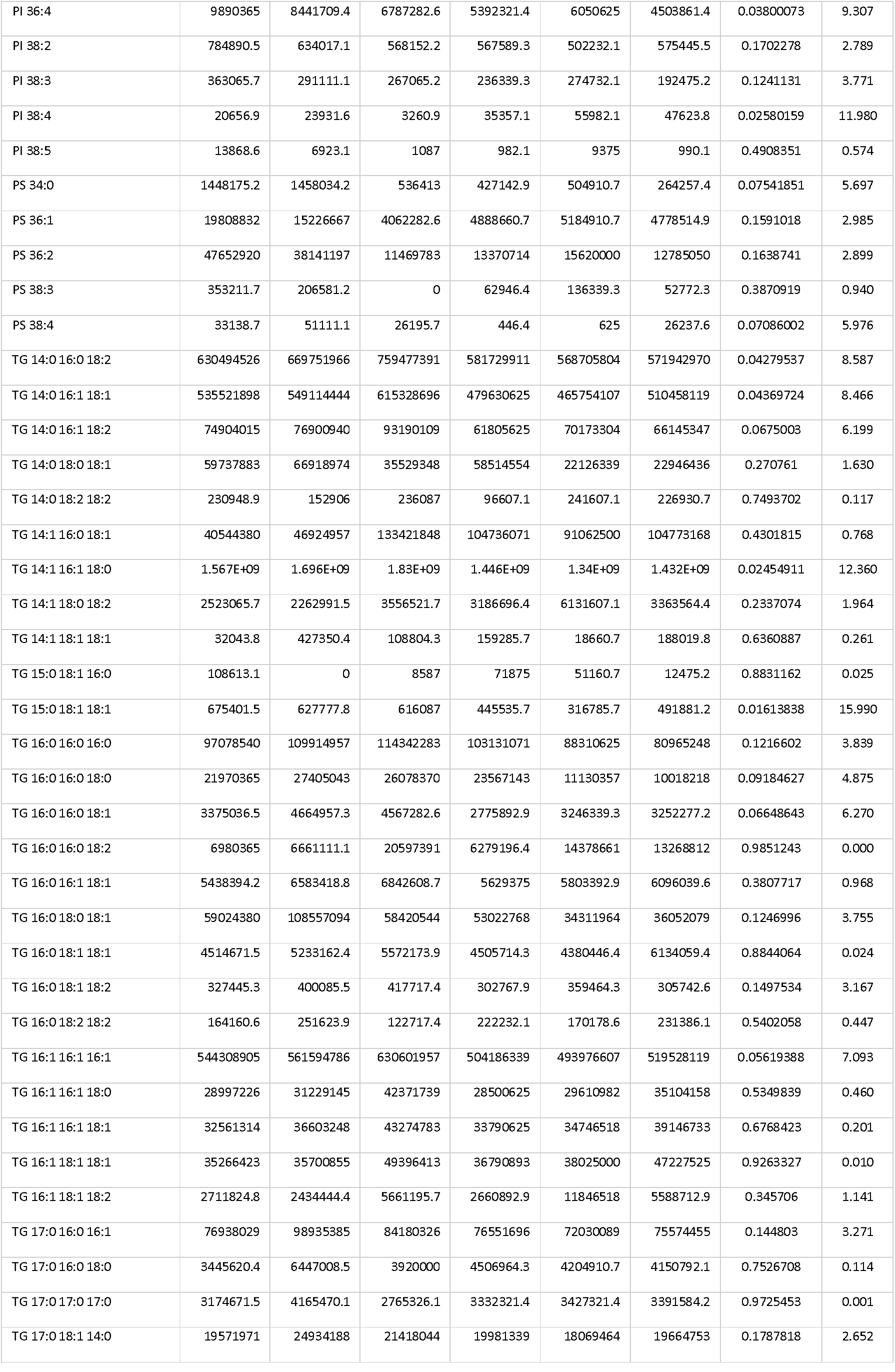
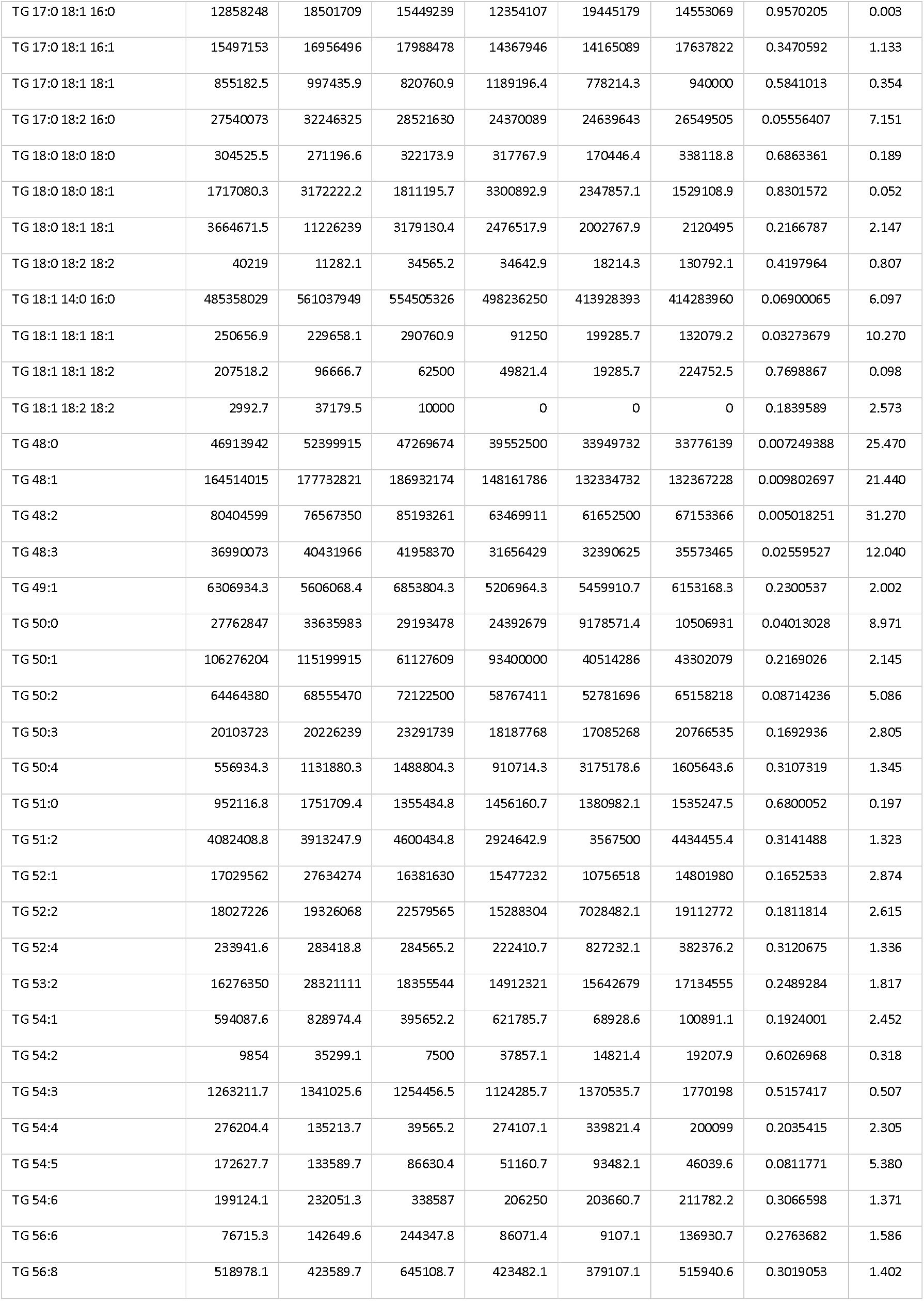
Impact of spinosad on the lipidomic profile. Lipidomic profile of larvae exposed to 2.5 ppm spinosad or control (equivalent dose of DMSO) for 2 hr as detected by LC-MS. Values are expressed as peak intensity area normalized to sample weight.

## Notes

### Competing Interest Statement

The authors have declared no competing interest.

